# Inflammation and bacteriophages affect DNA inversion states and functionality of the gut microbiota

**DOI:** 10.1101/2023.03.31.535104

**Authors:** Shaqed Carasso, Rawan Zaatry, Haitham Hajjo, Dana Kadosh-Kariti, Nadav Ben-Assa, Rawi Naddaf, Noa Mandelbaum, Sigal Pressman, Yehuda Chowers, Tal Gefen, Kate L. Jeffrey, Juan Jofre, Michael J. Coyne, Laurie E. Comstock, Itai Sharon, Naama Geva-Zatorsky

## Abstract

Reversible genomic DNA-inversions control expression of numerous bacterial molecules in the human gut, but how this relates to disease remains uncertain. By analyzing metagenomic samples from six human Inflammatory Bowel Disease cohorts combined with mice experimentation, we identified multiple invertible regions where a particular orientation was correlated with disease. These include the promoter of the anti-inflammatory polysaccharide-A (PSA) of *Bacteroides fragilis*, which is mostly oriented ‘OFF’ during inflammation but is present in the ‘ON’ orientation when inflammation is resolved. We further detected increased abundances of *B. fragilis*-associated bacteriophages in patients with the PSA ‘OFF’ orientation, and a significant reduction in the frequency of the ‘ON’ orientation, in the presence of the *B. fragilis*-associated bacteriophage, thereby altering the bacterial induced immune modulation. Altogether, we reveal dynamic and reversible bacterial phase-variations driven both by bacteriophages and the host inflammatory state, signifying bacterial functional plasticity during inflammation and opening future research avenues.

## Introduction

Phase variation is the process by which bacteria undergo reversible alterations in specific loci of their genome, resulting in ON-OFF expression of genes^1–4^. In Bacteroidales, the dominant order of bacteria in the human gut, phase variation is highly prevalent and is largely mediated by inversions of DNA segments between inverted repeats. These inversions often involve promoter regions dictating transcription initiation of genes or operons functioning as ‘ON’\‘OFF’ switches^5^. In addition, DNA inversions can occur so that new genes are brought from an inactive to a transcriptionally active site by re-orientation or recombination of genomic “shufflons”— mobile genetic elements that facilitate rearrangements, serving as dynamic tools for altering the expressed gene^6–8^. Analysis of the orientations of bacterial invertible regions in various host disease states can provide new insights into bacterial adaptation and functional contributions to the disease pathogenesis or its resolution. Phase variation in the gut Bacteroidales often modulates production of components presented on the bacterial surface^9^ dictating which surface molecules interact, for example, with neighboring microbes (e.g. bacteria, bacteriophages) or with the host.

*Bacteroides fragilis*, a common resident of the human gut, modulates its surface by the phase variable synthesis of its capsular polysaccharides (PS, denoted PSA-PSH). The biosynthesis loci of seven of its eight polysaccharides have invertible promoters that are orientated either ‘ON’ or ‘OFF’ in respect to the downstream PS biosynthesis operon^5^. Studies have shown that the *B. fragilis* polysaccharide A (PSA) modulates the host immune system by inducing regulatory T cells (Tregs) and secretion of the anti-inflammatory cytokine interleukin (IL)-10^10,11^. Moreover, PSA was shown to confer protection against experimental colitis^10,12–14^, and thus is regarded as an anti-inflammatory polysaccharide.

Bacteriophages, viruses that specifically target and infect bacteria, can influence the composition of bacterial populations and potentially affect their functionality. Several studies have highlighted the dynamic relationship between Bacteroidales and phages. Campbell et al.^15^ found that infection of *Bacteroides vulgatus* with temperate bacteriophage BV01 results in a repression of bile salt hydrolase activity – changing the bacteria transcriptional profile depending on the integration of the phages into their genome. Porter *et al.*^8^ and Hryckowian *et al.*^16^ demonstrated that some capsular PS and other outer surface molecules can affect the susceptibility of *Bacteroides thetaiotaomicron*, to specific phages. Consistent with that, Shkoporov *et al.*^17^ revealed that phase variation of capsular PS in *Bacteroides intestinalis* influences the infection of CrAss-like phages, establishing a dynamic equilibrium for their prolonged coexistence in the gut. These studies suggest that the interactions between phages and *Bacteroides* may affect the host through changes in bacterial functionality.

Ulcerative colitis (UC) and Crohn’s disease (CD) are multifactorial inflammatory bowel diseases (IBD), characterized by a compromised mucosal barrier, inappropriate immune activation, and mislocalization of the gut microbiota^18–21^. As such, IBDs have emerged as some of the most studied microbiota-linked diseases^22^ and an interesting setting for studying gut bacterial functions, their potential effects on the host, and bacteria-phage interactions.

Here we present an analysis of invertible DNA orientations in the gut microbiota of IBD patients from multiple cohorts and test the hypothesis that inflammation and bacteriophages mediate these differential orientations. Our combined analysis of IBD patient metagenomes and experimental mouse models, focusing on Bacteriodales species, reveals alterations in the orientations of multiple DNA invertible regions during gut inflammation with potential to modulate the host immune system. Importantly, we show that filtered fecal extracts of IBD patients alter the orientation of the PSA promoter of *B. fragilis*. We show that bacteriophages are correlated with the PSA ‘OFF’ state, and that a specific lytic bacteriophage of *B. fragilis* increases the PSA promoter ‘OFF’ population with concurrent decline of host colonic Treg cells. These findings reveal a dynamic interplay between gut inflammation, bacteriophages, and bacterial phase variation, with potential implications for the diagnosis and treatment of IBD.

## Results

### IBD is correlated with quantitative differences in the orientations of multiple Bacteroidales invertible DNA regions compared to healthy controls

As a means to determine if Bacteroidales phase variable molecules may be differentially produced in IBD patients compared to controls, we used the PhaseFinder^9^ software to identify and analyze the relative orientations of DNA inversions in the gut metagenomes of cohorts of IBD patients and control subjects **[Table 1]**. In brief, we first detected putative invertible regions in Bacteroidales genomes by identifying inverted repeats, and then created a database containing their forward and reverse orientations (compared to the submitted genome sequences). Metagenomic sequences from publicly available datasets were then aligned to the database, resulting in the ratio of the orientations of each invertible region compared to their orientations in the published genome sequence. Our initial analysis of 39 sequenced genomes including 36 human gut Bacteroidales species identified 311 invertible regions. Of these 311 invertible regions, there were 147 that were statistically different in terms of their orientations in IBD patients compared to healthy controls. These included invertible regions from 25 of the reference bacterial genomes **[Table S1]**. These regions included both invertible promoters and intragenic regions of diverse genes, spanning from regulatory genes to those encoding genes for the production of outer surface molecules. **Table S1** details the locations of the inverted repeats (IR) sequences and genes flanking the invertible regions **[Figure 1A]**. The regions in which the orientations differed most significantly between IBD patients and controls were within or in proximity to SusC/SusD-like outer membrane transport systems^23^ and to *upxY* genes, which, in most cases, represent the first gene in the biosynthesis loci of capsular polysaccharides (PS) **[Figure 1B]**. Four of the five invertible capsular polysaccharide promoters of *Bacteroides thetaiotaomicron* were differentially oriented between the groups, and three of the seven invertible PS promoters of *B. fragilis* were differentially oriented, including the anti-inflammatory PSA promoter being differentially oriented between healthy and UC patients. The PSA promoter showed a higher percentage (71%) of reverse oriented reads in IBD patients, compared to 56.2% in the healthy controls **[Figure 1C]**. In the reference genome of *B. fragilis* NCTC 9343, the PSA promoter is in its ‘ON’ orientation, hence, the reverse orientation, found in IBD patients, represents the PSA promoter’s ‘OFF’ orientation. We further identified differential promoter orientations in healthy and IBD cohorts for PULs and polysaccharides promoters, such as *Bacteroides thetaiotaomicro*n VPI-5482 **[Figure 1D]**, and *Phocaeicola dorei*, a bacteria shown to be present in healthy subjects^24,25^ and correlated with disease activity in UC^26^ **[Figure 1E]**.

**Table 1:** Metagenomic datasets included in this study.

| Dataset | N<br>(Samples) | CD | UC | Non-IBD | Country | Notes |  |
| --- | --- | --- | --- | --- | --- | --- | --- |
| IBDMDB | 1283 | 574 | 349 | 360 | United States | Longitudinal | Ref<sup>27</sup> |
| HMP (3 phases) | 320 |  |  | 320 | United States |  | Ref<sup>28</sup> |
| MetaHit | 122 | 4 | 21 | 97 | Denmark and Spain |  | Ref<sup>29</sup> |
| 1000IBD | 331 | 205 | 126 |  | the Netherlands |  | Ref<sup>30</sup> |
| GeversD_2014 | 50 | 36 |  | 14 | United States and Canada |  | Ref<sup>31</sup> |
| LewisJD_2015 | 303 | 303 |  |  | United States and Canada | Children with Crohn's Disease Longitudinal, treated | Ref<sup>32</sup> |
| Total | 2409 | 1122 | 496 | 791 |  |  |  |

**Figure 1:**
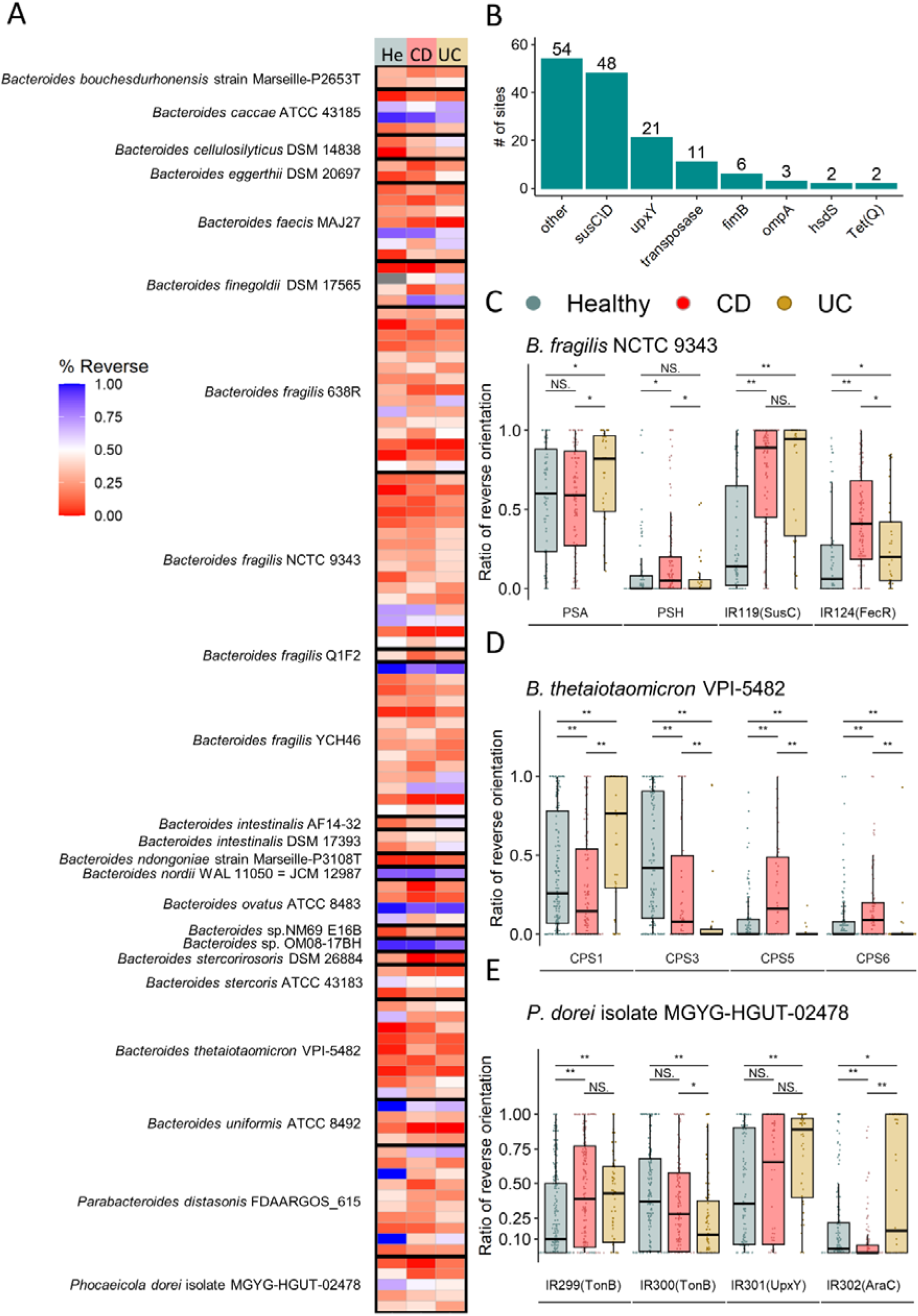
Bacteroides species exhibit differential orientations of invertible regions during health and disease. **A**. Selected significantly differentially oriented invertible DNA regions (inverted repeats (IR) segments, see **Table S1**) (Wilcoxon rank sum test, FDR p<0.05) in at least one comparison between Healthy, CD (Crohn’s Disease) and UC (Ulcerative Colitis). Blue indicates the Forward orientation and red represents the Reverse orientation in comparison to the reference genome. **B.** Prevalence of functional genes in proximity to invertible DNA regions significantly different between healthy individuals and IBD patients. **C.** Differentially oriented invertible DNA regions, in *B. fragilis* NCTC 9343. PSA: Polysaccharide A, PSH: Polysaccharide H. **D.** Differentially oriented invertible DNA regions in *B. thetaiotaomicron* VPI-5482. CPS: Capsular Polysaccharide. **E.** Differentially oriented invertible DNA regions in *Phocaeicola dorei* MGYG-HGUT-02478. (Wilcoxon rank sum test, \**p* < 0.05; \*\**p* < 0.01)

**Figure S1:**
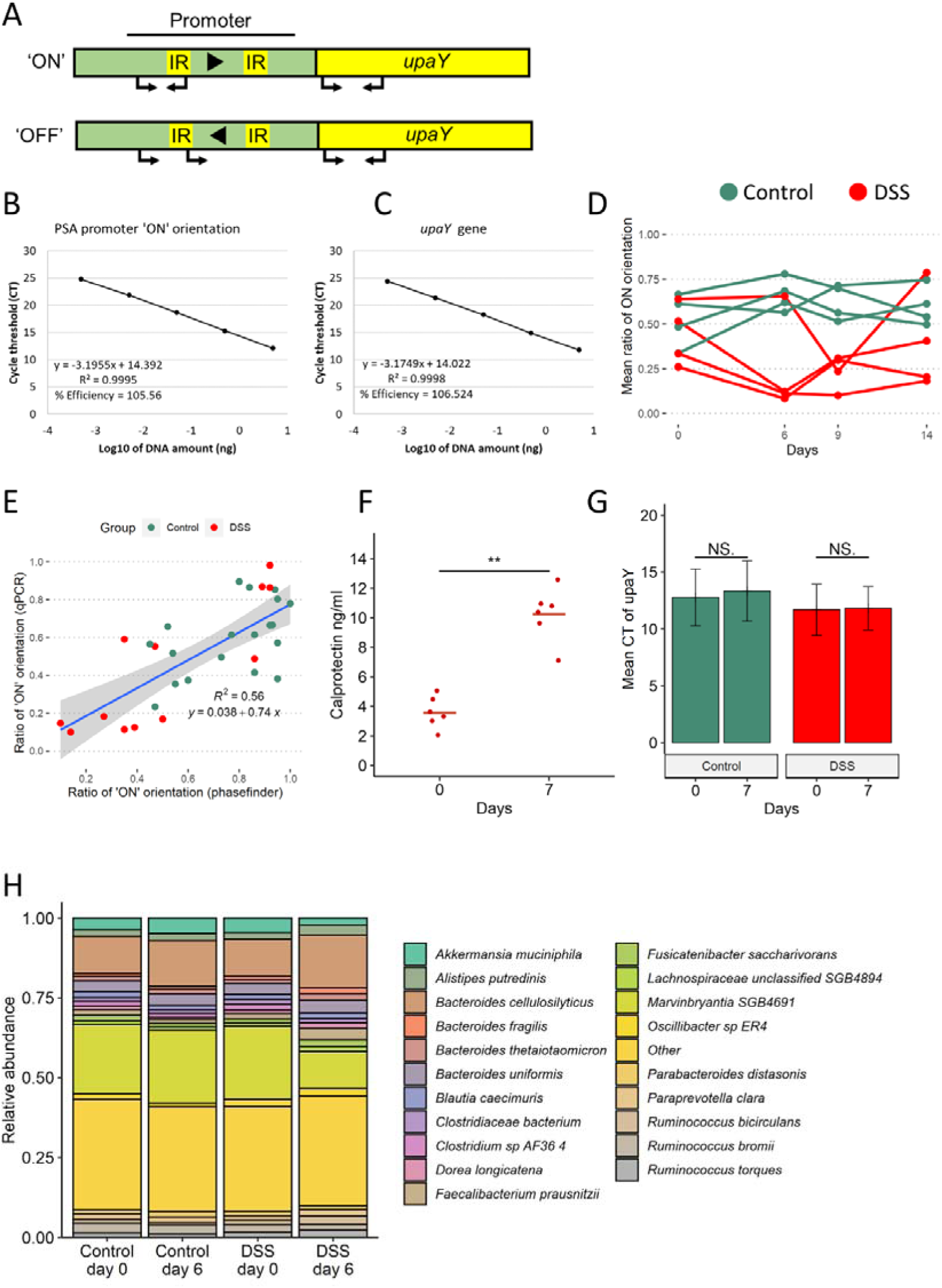
Relative orientation of the PSA promoter of *B. fragilis* is affected by inflammation. **A**. Schematics of the qPCR assay used to assess the ratio of PSA ‘ON’ promoter orientation. Primers (arrows) were directed upstream and within the PSA promoter invertible region (IR: inverted repeats) targeting only the ‘ON’ oriented promoters. Another set of primers was directed at the *upaY* and used to normalize the results to the number of genomes in a sample. **B. & C.** Primer efficiencies for the two sets of primers. Standard curves were calculated for each primer set from the Cycle thresholds (CTs) and the log of the initial DNA concentrations. Each sample represents the mean CT of 3 technical repeats. **D.** Mean ratio of *B. fragilis* PSA’s promoter ‘ON’ orientation measured by qPCR for different experiments on different days (n=8-12 in each time point). Green: Control group, Red: DSS treated mice. **E.** Scatter plot of the ratio of *B. fragilis* PSA’s promoter ‘ON’ orientation measured by qPCR and by the PhaseFinder tool. Each point represents a single sample, colored according to group and timepoints: green: Control group, red: DSS treated mice. Blue line represents linear regression (gray area represents 95% confidence intervals). **F.** Calprotectin levels (ng/ml) measured in gnotobiotic mice colonized with *B. fragilis* and treated with DSS on different days of the experiment. Line represents the mean. (Wilcoxon rank sum test, \*\**p* < 0.01) **G.** Mean qPCR cycle threshold (CTs) of *upaY* gene DNA, representing the abundance of *B. fragilis* in gnotobiotic mice monocolonized with *B. fragilis* of different groups and timepoints. Bars represent the means of the CTs in each group and timepoint. Error bars represent the standard error. Green: Control group, Red: DSS treated mice. (Wilcoxon rank sum test, *p*> 0.05). **H.** Stacked bar plots representing the relative abundance of the 20 most abundant bacterial species in the humanized mice in each experimental group in days 0 and 6.

### Differential promoter orientation states are induced by inflammation in a dynamic and reversible manner

To assess the dynamics of the *B. fragilis* PSA promoter orientation under inflammatory conditions, we designed a longitudinal experimental mouse model during which we analyzed the PSA promoter orientation over time. Germ-free (GF) mice were colonized with the gut microbiota from a healthy human microbiota, and subsequently spiked with *B. fragilis* NCTC 9343, termed “humanized” mice. Progeny of these mice were used for the experiments (n=8-12 mice per experimental group, DSS vs. control). Experimental colitis was induced by adding 3% dextran sodium sulfate (DSS) to the drinking water^33^ for 9 days after which DSS was replaced with water until day 14 **[Figure 2A]**. Inflammation was monitored by the levels of calprotectin in the stool **[Figure 2B]**, a commonly used biomarker for inflammation, and by mouse weight loss **[Figure 2C]**. Stool was collected before, during and after DSS treatment to analyze orientations of specific invertible DNA regions, using quantitative PCR (qPCR) to observe the state of the PSA promoter **[Figure S1A-C]**. The orientation of the *B. fragilis* PSA promoter is variable to some extent in both the control and DSS-treated mice and was significantly altered on day 9 in the DSS-treated mice along with the inflammatory state **[Figure 2B-D]**. At the beginning of the experiment, ∼45% of the population had the promoter in the ‘ON’ orientation **[Figure 2D]**. A significant decline in these percentages was observed six days after DSS was introduced (down to an average of ∼26%) and remained at around ∼24% on average on day nine. By day 14, when the mice were starting to gain weight and the calprotectin levels were decreasing **[Figure 2B,2C]**, the bacterial population returned to an average of 40% ‘ON’, similar to the beginning of the experiment, and to the control group, which did not receive DSS (mean of 58% of the *B. fragilis* population ‘ON’) **[Figure 2D]**.

**Figure 2:**
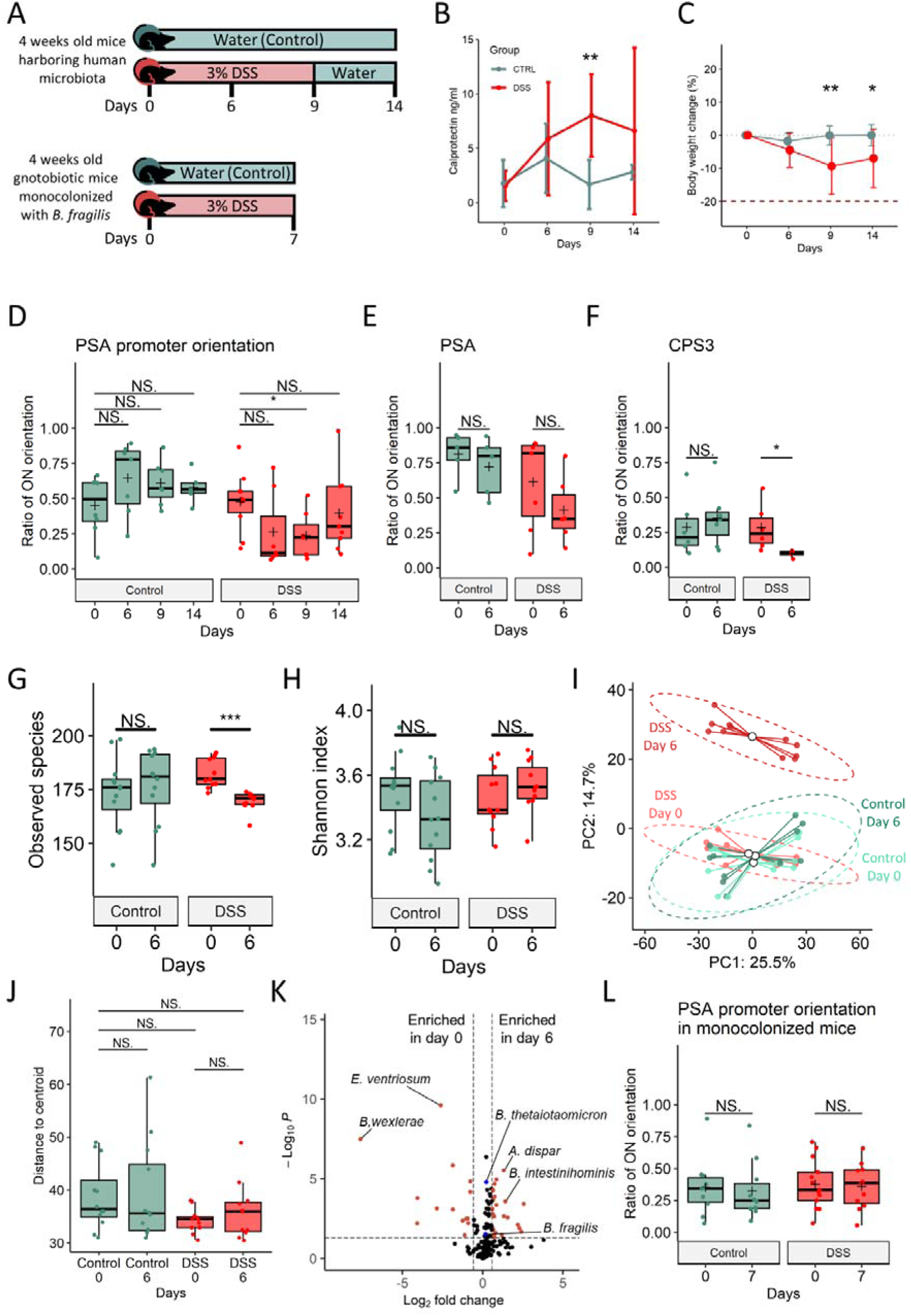
Relative orientation of the PSA promoter of *B. fragilis* is affected by inflammation. **A**. Illustration of murine model of inflammation. “Humanized’’ mice harboring human microbiota spiked with *B. fragilis* NCTC 9343, and monocolonized with *B. fragilis* NCTC 9343 were exposed to 3% DSS (Day=0 in the illustration) at four weeks of age; **B.** Calprotectin levels (ng/ml) measured in different days of the experiment. Lines represent the standard deviations. Green: Control group, Red: DSS treated mice. **C.** The body weight change of mice measured in different days of the experiment. Lines represent the standard deviations. Green: Control group, Red: DSS treated mice. **D.** Ratio of *B. fragilis* PSA’s promoter ‘ON’ orientation measured by qPCR on different days of the experiment (n=8 in each time point). Data represents the mean (cross), median (line in box), IQR (box), and minimum/maximum (whiskers). (Wilcoxon rank sum test, \**p* < 0.05; \*\**p* < 0.01). Green: Control group, Red: DSS treated mice. **E.** Ratio of *B. fragilis* PSA’s promoter reverse orientation measured by the PhaseFinder tool, in different days of the experiment. Data represent the median (line in box), IQR (box), and minimum/maximum (whiskers). (Wilcoxon rank sum test, \**p* <0 .05; \*\**p* <0 .01). Green: Control group, Red: DSS treated mice. **F.** Ratio of *B. thetaiotaomicron* CPS3’s promoter reverse orientation measured by the PhaseFinder tool on different days of the experiment. Data represent the median (line in box), IQR (box), and minimum/maximum (whiskers). (Wilcoxon rank sum test, \**p* < 0.05; \*\**p* < 0.01). Green: Control group, Red: DSS treated mice. **G.** Alpha diversity (observed species) between groups and timepoints. Data represent the median (line in box), IQR (box), and minimum/maximum (whiskers). (Wilcoxon rank sum test, \**p* < 0.05; \*\**p* < 0.01, \*\*\**p*<0.001). Green: Control group, Red: DSS treated mice. **H.** Alpha diversity (Shannon index) between groups and timepoints. Data represent the median (line in box), IQR (box), and minimum/maximum (whiskers). (Wilcoxon rank sum test, \**p* <0 .05; \*\**p* < 0.01, \*\*\**p*<0.001). Green: Control group, Red: DSS treated mice. **I.** Beta diversity (Aitchison distance) between groups and timepoints. Principal Coordinates Analysis (PCoA) of Aitchison distances between bacterial communities of different groups and timepoints. Each point represents a single sample, colored according to group and timepoints: light Green: Control on day 0, Green: control on day 6, light red: DSS treated on day 0, red: DSS treated on day 6. The mean (centroid) of samples in each group is indicated with a blank circle. Ellipses represent 0.95 confidence intervals of each group. **J.** Beta-dispersion values (Aitchison distances from the centroid) between groups and timepoints. Data represent the median (line in box), IQR (box), and minimum/maximum (whiskers). (betadisper, *p*> 0.05). **K.** Differential bacterial abundance between DSS treated mice in day 0 and day 6 detected by the Maaslin2 algorithm. Red dots indicate differentially abundant bacteria that were determined by adjusted *P* value < 0.05 and log_2_ fold change >1 and <-1, respectively. Blue dots indicate *B. fragilis* and *B. thetaiotaomicron*. **L.** Ratio of *B. fragilis* PSA’s promoter ‘ON’ orientation measured by qPCR in different days in gnotobiotic mice monocolonized with *B. fragilis* NCTC 9343. Data represent the median (line in box), IQR (box), and minimum/maximum (whiskers). (Wilcoxon rank sum test, \**p* <0 .05; \*\**p* < 0.01). Green: Control group, Red: DSS treated mice.

Since the frequency of *B. fragilis* with the PSA promoter oriented ‘ON’ declined in most of the inflamed mice **[Figure S1D]**, we performed metagenomic sequencing from stool collected on days zero and six for analysis of orientations of invertible regions in other Bacteroidales species and genomic regions **[Table S2]**.

This analysis validated the decline in the frequency of the ‘ON’ orientation of the *B. fragilis* PSA promoter **[Figure 2E]**, aligned with the qPCR results **[Figure S1E]**, and agreeing with the human metagenomic analysis **[Figure 1C]**. In addition, we identified a decline in the frequency of the ‘ON’ orientation of the CPS3 promoter of *B. thetaiotaomicron* during inflammation in mice **[Figure 2F]**, consistent with the results of the human metagenomic analysis where it also showed lower ratios of the ‘ON’ orientation, equivalent to reverse oriented reads in comparison to the reference genome **[Figure 1D]**.

The bacterial composition of the gut microbiota in control mice was stable throughout the experiment but varied in the DSS-treated mice. Bacterial richness (alpha-diversity, observed species) declined during inflammatory conditions **[Figure 2G]**, while evenness (alpha-diversity, Shannon index) remained constant **[Figure 2H]**. The beta-diversity (Aitchison distance) was significantly altered **[Figure 2I],** while the beta-dispersion (Aitchison distance) was not significantly altered **[Figure 2J]**.

Inflamed mice showed a decrease in relative abundances of *Blautia* species and *Eubacterium ventriosum*, with a concomitant increase in *Barnesiella intestinihominis* and *Alistipes dispar* **[Figure 2K, Table S2]**. The relative abundance of *B. fragilis*, which exhibited differential orientations of invertible DNA regions, was not significantly altered during inflammation. **[Figure 2K]**.

To better assess the role of inflammation on the orientation of the *B. fragilis* PSA promoter we repeated the experiment using gnotobiotic mice mono-colonized with *B. fragilis* NCTC 9343 (n=8-12 mice per experimental group, DSS vs. control), excluding influences of altered microbiota **[Figure 2A, S1F]**. The PSA promoter orientation was not altered in the monocolonized mice **[Figure 2L]**, along with the bacterial abundances **[FIGURE S1G]** over the course of the experiment (with or without DSS), suggesting a role of the microbiota in driving the ‘OFF’ orientation of the PSA promoter during inflammation.

### The inflammatory milieu of a complex human microbiota results in the PSA promoter “OFF” orientation

We next sought to examine factors of the complex microbiota that result in a decreased percentage of the *B. fragilis* population with the PSA promoter orientated ‘ON’ during inflammation. To do so, we exposed *B. fragilis* NCTC 9343 to fecal filtrates from IBD patients. CD and UC patients were recruited from the Rambam Health Care Campus (RHCC). Fecal samples were collected before and after treatment (Infliximab (HR) or Humira (HuR) therapy, 4 and 7 patients respectively). Both treatments are antibodies targeted against tumor necrosis factor-α (TNF-α), an inflammatory cytokine increased in IBD patients. *B. fragilis* NCTC 9343 was cultured in fecal filtrates to mid-log phase, at which time, DNA was extracted for qPCR analysis of the PSA promoter orientation **[Figure 3A]**. *B. fragilis* exposed to fecal filtrates of patients before treatment showed higher ratios of the population with the PSA promoter oriented ‘OFF’, while *B. fragilis* exposed to fecal filtrates after treatment showed higher ratios of the PSA promoter oriented ‘ON’ **[Figure 3B]**. This observation was in line with the fecal calprotectin concentrations which were high before treatment and decreased after treatment, inversely correlating with PSA promoter ‘ON’ orientation **[Figure 3C]**. These results demonstrate that the population of bacteria with the PSA promoter in each orientation varies in the inflamed and non-inflamed gut, with a higher percentage of the population in the ‘OFF’ orientation under inflammatory conditions, and a shift towards the ‘ON’ orientation following reduction in inflammation **[Figure 3B, 3C]**. Particularly, treatment responders had a significant correlation of PSA ‘ON’ with lower calprotectin levels, compared to the non-responders **[Figure S2A]**. This change could be due to induction of inversion under these conditions or to selection of a preferred promoter state.

**Figure 3:**
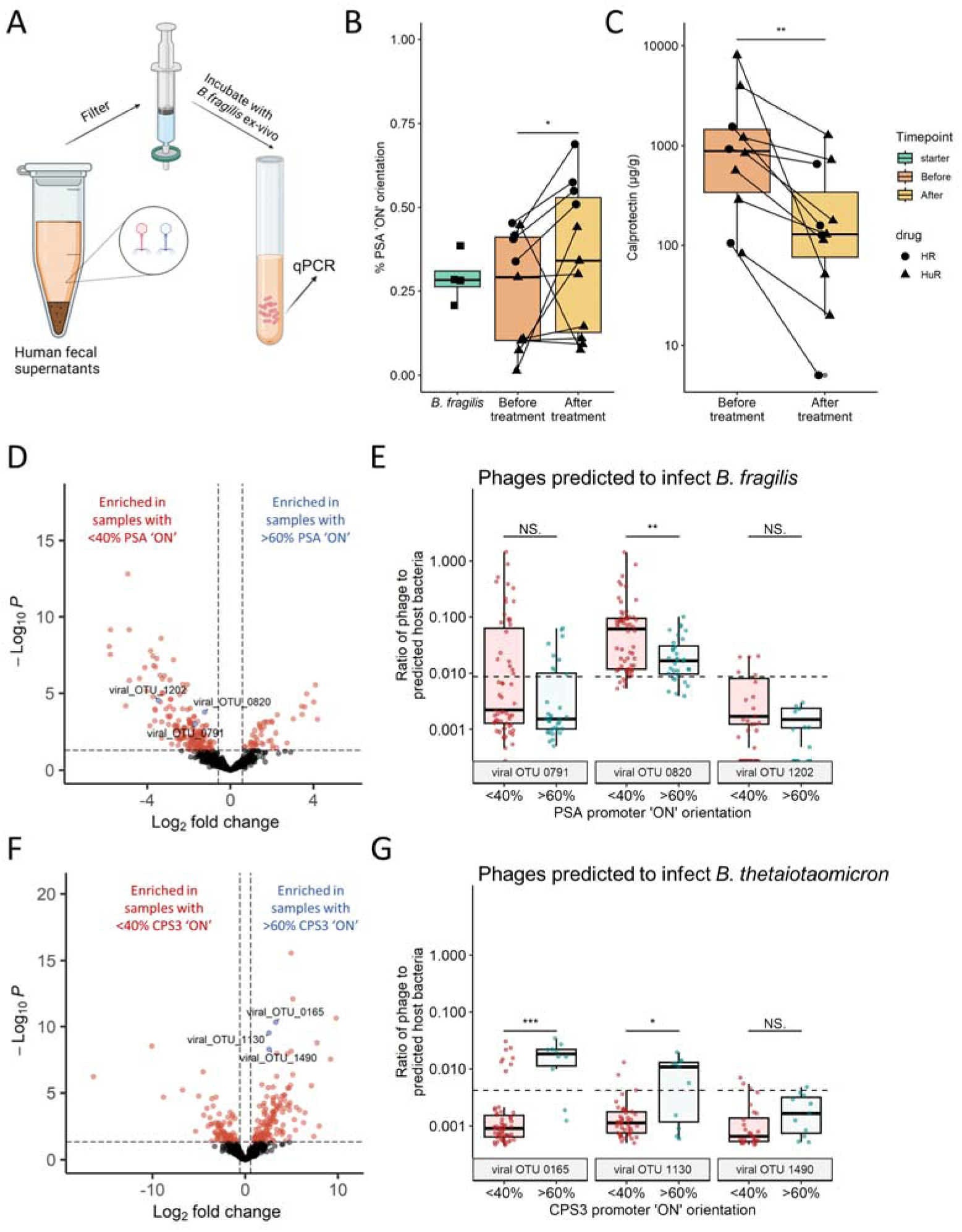
Evidence of phage correlation with *B. fragilis* polysaccharide A promoter orientation. **A**. Experimental design of culturing *B. fragilis* in patients’ fecal filtrates. **B.** Ratio of the ‘ON’ orientation of the PSA promoter of *B. fragilis,* measured by qPCR, after ex-vivo exposure to fecal filtrates of IBD patients before and after treatment with anti-TNF (4 patients on Infliximab (HR) and 7 patients on Humira (HuR)). Data represent the median (line in box), IQR (box), and minimum/maximum (whiskers). (One-sided Wilcoxon rank sum test, \**p* <0 .05). green: *B. fragilis* grown in a 1:1 ratio of M9 and PBS not exposed to fecal filtrates, orange: *B. fragilis* exposed to fecal filtrates of patients before anti-TNF treatment. yellow: *B. fragilis* exposed to fecal filtrates of patients after treatment. Each dot represents 3 individual experiments; lines connect experiments from the same patient; Shapes are determined by the patients’ treatments, circle: HR, triangle: HuR. **C.** Calprotectin levels (µg/g) measured in patients’ feces. Data represent the median (line in box), IQR (box), and minimum/maximum (whiskers). (One-sided Wilcoxon rank sum test, \**p* <0 .05). Dots represent samples; lines connect samples from the same patient; Shapes are determined by the patients’ treatments, circle: HR, triangle: HuR. **D.** Differential viral taxonomic units’ abundances, (from the IBDMDB cohort, count table from Nishiyama *et al.* (2020).), between samples with low ‘ON’ orientation of the PSA promoter (<40%) and high ‘ON’ orientation (>60%). Differentially abundant viral taxonomic units were detected by the DeSeq2 algorithm (Wald test, p < 0.01). Red dots indicate differentially abundant bacteria that were determined by P value < 0.01 and fold change >1.5 and <-1.5, respectively. **E.** Phage to host ratios of viral OTUs predicted to infect *B. fragilis* and detected in Figure 3D. Data represent the median (line in box), IQR (box), minimum/maximum (whiskers), dots represent individual samples. Dashed line represents the mean phage to host ratio of all phages predicted to infect *B. fragilis* in all samples (mean = 0.009). **F.** Phage to host abundances ratios of viral OTUs 0791 and 0820 against samples’ levels of PSA’s promoter ‘ON’ orientation (red: <40%, blue: >60%) analyzed from the IBDMDB cohort, based on the count table from Nishiyama *et al.* (2020). (Wilcoxon rank sum test, \**p* <0 .05; \*\**p* <0 .01) **G.** Differential viral taxonomic units’ abundances, (from the IBDMDB cohort, count table from Nishiyama *et al.* (2020).), between samples with low ‘ON’ orientation of the CPS3 promoter (<40%) and high ‘ON’ orientation (>60%). Differentially abundant viral taxonomic units were detected by the DeSeq2 algorithm (Wald test, p < 0.01). Red dots indicate differentially abundant bacteria that were determined by P value < 0.01 and fold change >1.5 and <-1.5, respectively. **H.** Phage to host abundances ratios of viral OTUs predicted to infect *B. thetaiotaomicron* and detected in Figure 3G. Data represent the median (line in box), IQR (box), minimum/maximum (whiskers), dots represent individual samples. Dashed line represents the mean phage to host ratio of all phages predicted to infect *B. thetaiotaomicron* in all samples (mean = 0.03).

**Figure S2:**
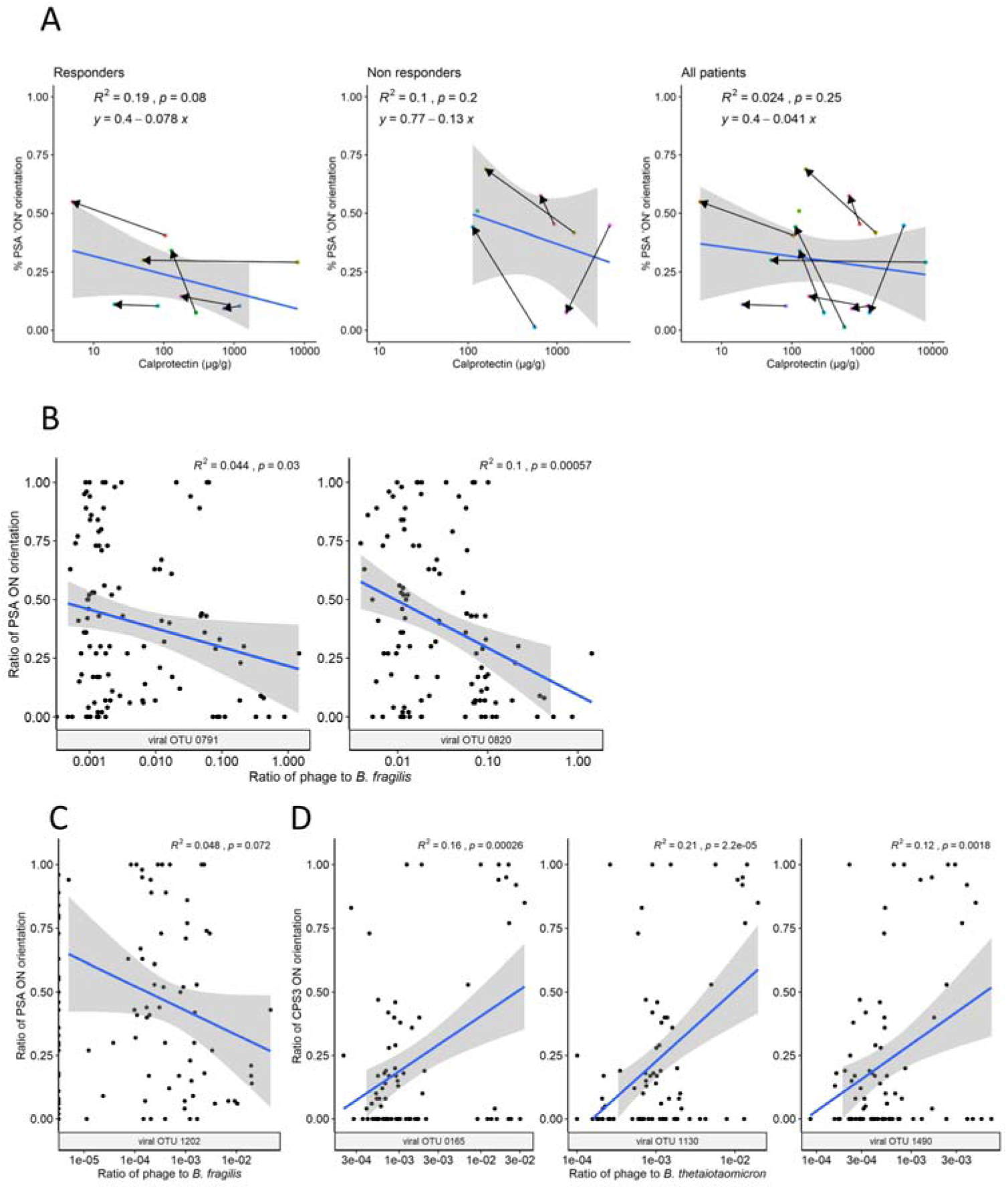
Phage correlation with *B. fragilis* polysaccharide A promoter orientation. **A**. Scatter plots of the ratio of *B. fragilis* PSA’s promoter ‘ON’ orientation measured by qPCR and fecal calprotectin levels (µg/g). Each point represents a single sample, colors indicate the same patient, arrows denote the change from samples before anti-TNF treatment to after. Blue line represents linear regression (gray area represents 95% confidence intervals). **B. & C.** Scatter plots of the ratio of *B. fragilis* PSA’s promoter ‘ON’ orientation measured by qPCR and phage to host ratios of viral OTUs predicted to infect *B. fragilis* in Nishiyama *et al.* (2020). Blue line represents linear regression (gray area represents 95% confidence intervals). **D.** Scatter plots of the ratio of *B. fragilis* PSA’s promoter ‘ON’ orientation measured by qPCR and phage to host ratios of viral OTUs predicted to infect *B. thetaiotaomicron* in Nishiyama *et al.* (2020). Blue line represents linear regression (gray area represents 95% confidence intervals).

### Investigating associations of phage with PSA promoter orientation

Fecal filtrates contain a mixture of bacterial and host metabolites, cytokines, antibodies, viruses, bacteriophages, and other components. Bacteriophages were shown to be associated with intestinal inflammation and IBD in several studies^34–38^, including Nishiyama *et al.*^39^ who characterized temperate bacteriophages and their bacterial hosts in the IBDMDB metagenomics database. Here, using phage sequences from the IBDMDB database, we compared the relative abundances of these phages between samples that displayed lower (<40%) or higher (>60%) ratios of the PSA promoter ‘ON’ orientation. Differential relative abundance analysis revealed 108 viral OTUs enriched in patients with lower ratios of the PSA promoter ‘ON’ orientation after FDR correction, compared to 24 viral OTUs enriched in patients with the higher ratios of the ‘ON’ orientation] **Figure 3D, Table S3**[. Three of the bacteriophages enriched in the lower ratios of the PSA promoter ‘ON’ orientation was predicted to infect *B. fragilis*^39^ – viral OTUs 0791, 0820, and 1202. To assess whether the relative abundances of the bacteriophages were accompanied with lower relative abundances of *B. fragilis*, the phage-to-host abundance ratio was calculated for each phage-*B. fragilis* pair. Two of the OTUs whose relative abundances were correlated to the PSA promoter ‘OFF’ orientation, viral OTUs 0791 and 0820, also showed higher phage-to-host abundance ratios **[Figure 3E, Figure S2]**. Intriguingly, these two viral OTUs were found to be more abundant in active CD and UC^39^.

We conducted the same analysis comparing samples with high to low ‘ON’ orientation ratios of the CPS3 promoter of *B. thetaiotaomicron*. Although there were differences in the virome compositions between these groups, no *B. thetaiotaomicron* associated bacteriophages were correlated with the ‘OFF’ orientation of the CPS3 promoter (i.e., the orientation, which we identified as associated with the disease) **[Figure 3F, Table S3]**. We found three *B. thetaiotaomicron* associated with viral OTUs – 0165, 1130, and 1490, slightly associated with the ‘ON’ orientation of the CPS3 promoter (the orientation that we identified as associated with healthy controls) **[Figure 3G, Figure S2]**. In addition, these bacteriophages displayed low phage to host ratios **[Figure 3G]**. To note, these OTUs were not previously associated with disease^39^.

### Bacteriophage exposure correlates with *B. fragilis* PSA promoter ‘OFF’ state

Since a higher abundance of bacteriophages were correlated with the PSA promoter in the ‘OFF’ orientation, we sought to study whether encounter with bacteriophage can be associated with altered orientation of the PSA promoter in *B. fragilis*. To this end, we isolated from sewage Barc2635 – a *B. fragilis* NCTC 9343 specific lytic bacteriophage **[Figure S3A]**, sequenced its genome, (deposited to GenBank, accession: MN078104), and characterized its morphology by electron microscopy **[Figure S3B].**

**Figure S3:**
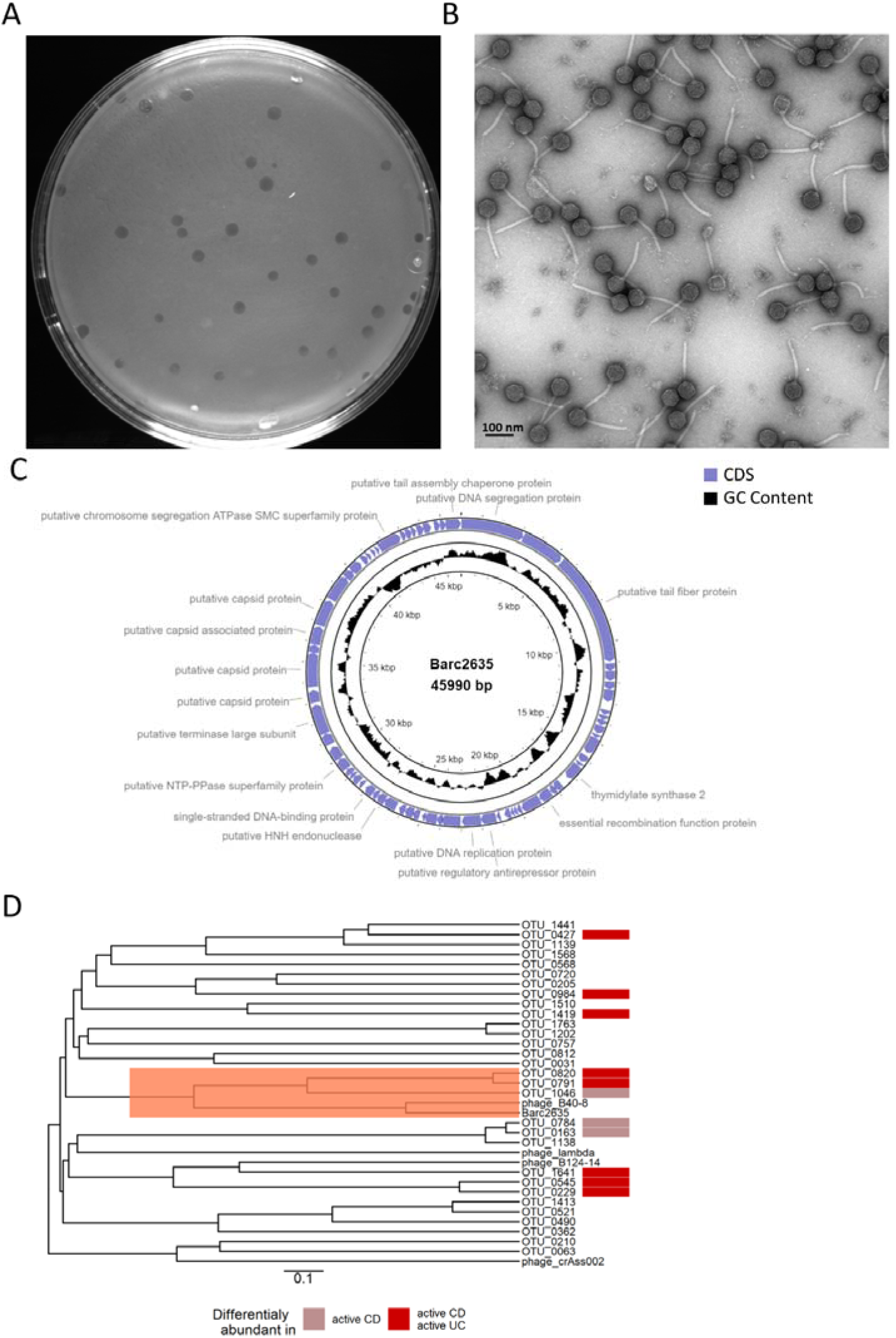
Characterization of Bacteriophage Barc2635. **A**. Barc2635 plaques on a lawn *B. fragilis* NCTC 9343. **B.** A representative transmission electron microscopy of Barc2635. **C.** Genome structure, GC content, and putative annotations of Barc2635. CDS: coding sequence. **D.** Phylogenetic tree based on the whole genome of viral OTUs identified as bacteriophages against *B. fragilis* as well as *Bacteroides* bacteriophages Barc2635, B40-8, B124-14, crAss002, and *Enterobacteria* phage lambda. MAFFT was used to perform multiple sequence alignment and the average-linkage method was used to construct the phylogenetic tree. Colors denote association between the viral OTUs abundances in Nishiyama *et al.* (2020), IBDMDB cohort, with active Crohn’s disease (light red) or with both active Ulcerative colitis and active Crohn’s disease (red).

Barc2635 is a lytic double-stranded DNA bacteriophage of 45,990 bp with a GC content of 38.9%, containing 67 putative CDS belonging to the tailed phages unified in the class Caudoviricete, formerly *Caudovirales*, **[Figure S3C]**. Interestingly, increased abundance of bacteriophages from this class were found to be correlated with IBD patients (UC and CD)^35^. We analyzed the sequence similarities of Barc2635, with the *B. fragilis* associated bacteriophages identified in the IBDMDB^39^ as well as from other studies^40–42^, and found that Barc2635 is most similar to a cluster of bacteriophages, including viral OTUs 0791 and 0820 which were associated with active disease^39^ **[Figure 4A, S3D]**.

**Figure 4:**
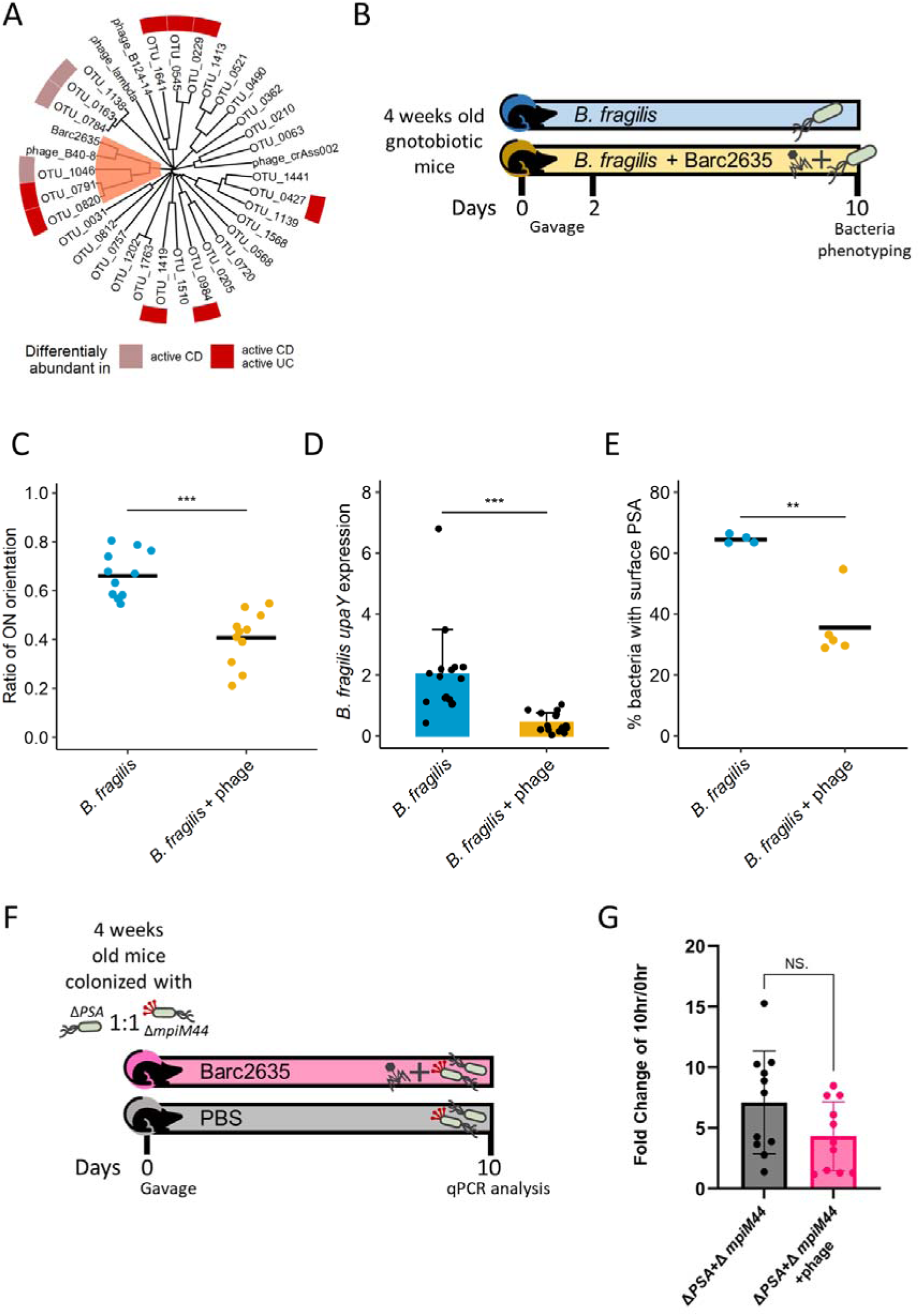
Phage exposure alters expression of the PSA locus and surface PSA. **A**. Phylogenetic tree based on the whole genome of viral OTUs identified as bacteriophages against *B. fragilis* as well as *Bacteroides* bacteriophages Barc2635, B40-8, B124-14, crAss002, and *Enterobacteria* phage lambda. MAFFT was used to perform multiple sequence alignment and the average-linkage method was used to construct the phylogenetic tree. Colors denote association between the viral OTUs abundances in Nishiyama *et al.* (2020), IBDMDB cohort, with active Crohn’s disease (light red) or with both active Ulcerative colitis and active Crohn’s disease (red). **B.** Experimental design of *in vivo* experiments. **C.** Ratio of *B. fragilis* PSA’s promoter ‘ON’ orientation measured on day 10 by qPCR in fecal samples of gnotobiotic mice monocolonized with *B. fragilis* with and without the Barc2635 bacteriophage. Horizontal lines represent the mean. Blue: Control group, Red: Barc2635 treated mice. Each dot represents a mouse. (Mann-Whitney test, \*\*\**p* < 0.001). **D.** Expression levels of *upaY* in ceca of gnotobiotic mice monocolonized with *B. fragilis* with and without the Barc2635 bacteriophage on day 10. Levels are shown as 2^(–ΔCT) with *rpsL* as a reference gene. Each dot represents a mouse. (Mann-Whitney test, \*\*\*\**P*<0.0001). **E.** PSA presence on the surface of *B. fragilis* exposed to the Barc2635 bacteriophage, detected by anti-PSA antibodies on day 10. Bacteria were analyzed by flow cytometry for the expression of PSA, using Rabbit anti PSA antibodies. Each dot represents a mouse. **F.** Experimental design of *in vivo* competition experiments. **G.** Fold change of Δpsa\ΔmpiM44 at Tp10 and Tp0 in the Barc2635 bacteriophage treated group (pink) compared to Δpsa\ΔmpiM44 at Tp10 and Tp0 in the control group (black). Each dot represents a mouse. (Mann-Whitney test, p>0.05).

**Figure S4:**
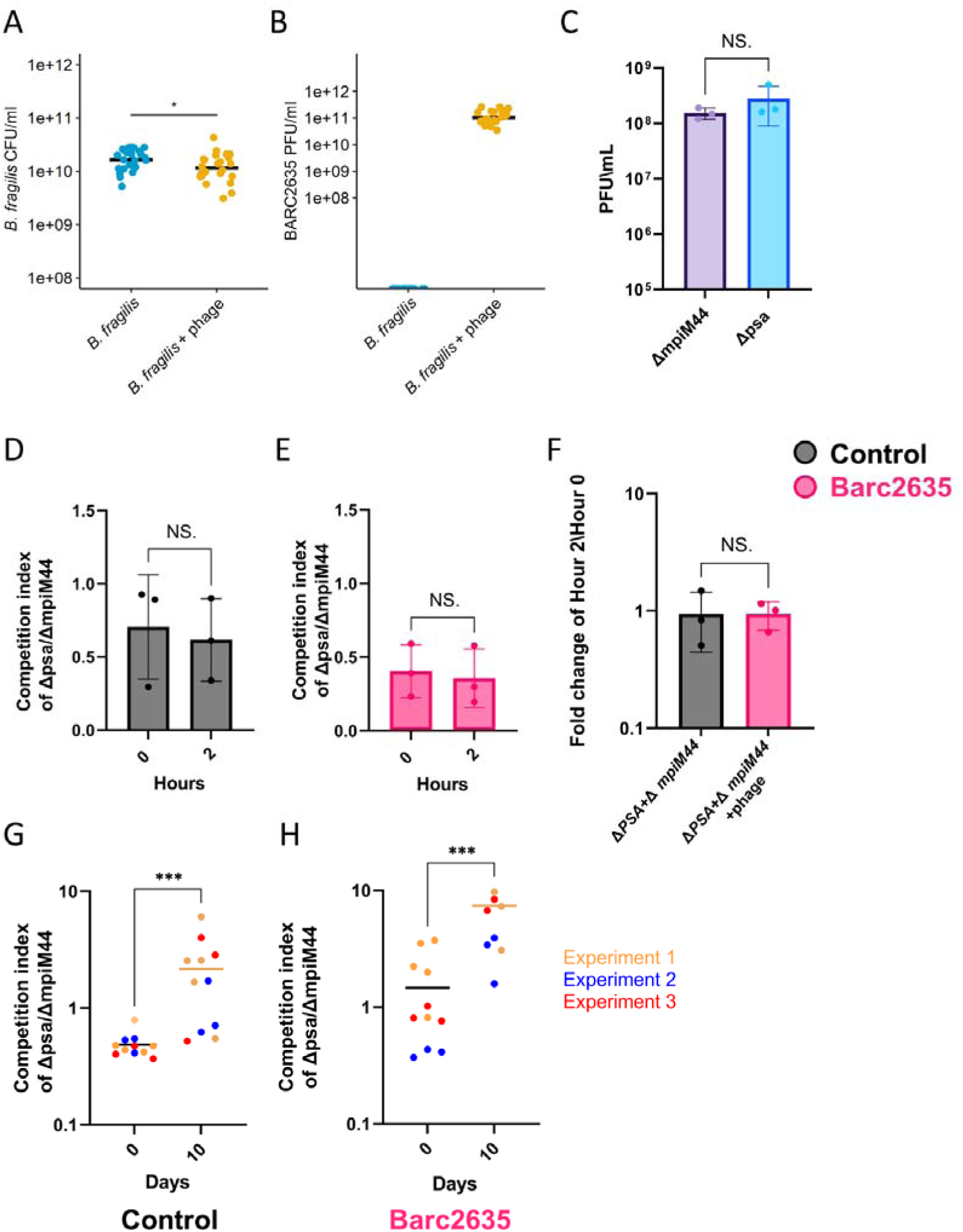
Barc2635 infects both Δmpi and Δpsa in the same manner in vivo. **A**. CFUs of *B. fragilis* in fecal samples of mice monocolonized with *B. fragilis* (blue) or colonized with *B. fragilis* + phage (yellow) measured on day 10 (Wilcoxon rank sum test, *p < 0.05). **B**. PFUs of Barc2635 in fecal samples of mice monocolonized with *B. fragilis* (blue) or colonized with *B. fragilis* + phage (yellow) measured on day 10. **C**. Barc2635 plaque forming units of ΔmpiM44 (purple) compared to Δpsa (blue). Each dot represents a different experiment. (Mann-Whitney test, p>0.05). **D**. Competition index of Δpsa\ΔmpiM44 in the control group at the beginning of the experiment (0 horse) compared to 2 hours, measured by qPCR. Each dot represents a different experiment (Mann-Whitney test, p>0.05). E. Competition index of Δpsa\ΔmpiM44 in Barc2635 treated group at the beginning of the experiment (0 horse) compared to 2 hours, measured by qPCR. (Mann-Whitney test, p>0.05) **F.** Fold change of Δpsa\ΔmpiM44 at the beginning of the experiment (0 horse) compared to 2 hours in the Barc2635 bacteriophage treated group (pink) compared to Δpsa\ΔmpiM44 at Tp2 and Tp0 in the control group (black). Each dot represents a different experiment. (Mann-Whitney test, p>0.05). **G.** Competition index of Δpsa\ΔmpiM44 in the control group on day 0 compared to day 10, measured by qPCR. Each dot represents a mouse (Mann-Whitney test, *\*\*\*p<0.001*). **H.** Competition index of Δpsa\ΔmpiM44 in the treated group on day 0 compared to day 10, measured by qPCR. Each dot represents a mouse (Mann-Whitney test, *\*\*\*p<0.001*)

To test whether exposure to this bacteriophage leads to changes in relative orientations of the PSA promoter of the *B. fragilis* population, we monocolonized GF mice with *B. fragilis* NCTC 9343 in the presence or absence of Barc2635 and analyzed the PSA promoter DNA inversion state **[Figure 4B]**. In response to bacteriophage, a higher percentage of the population had the PSA promoter in the ‘OFF’ state **[Figure 4C].** To further analyze the consequence of PSA promoter inversion we measured the mRNA expression of the first gene in the PSA biosynthesis locus – *upaY*^43^, which requires the PSA promoter to be oriented ‘ON’. **Figure 4D** shows a significant reduction in *upaY* expression in mice containing Barc2635 in comparison to mice with bacteria alone. To directly measure the levels of PSA on the surface of the *B. fragilis* population, we used specific antibodies and monitored surface PSA by flow cytometry. The percentage of bacterial cells synthesizing PSA from mice with bacteriophage Barc2635 was significantly lower in comparison to bacterial cells from mice without the bacteriophage **[Figure 4E]**, in agreement with the *upaY* expression levels and the PSA promoter orientation. In the two groups of mice (i.e., *B. fragilis* alone and *B. fragilis* with Barc2635), the CFU levels of *B. fragilis* were different by 0.5 a log **[Supp. Figure S4A]**, and no phages were detected in the *B. fragilis* monocolonized group (i.e., without bacteriophage) **[Supp. Figure S4B]**.

### Barc2635 infects both ΔPSA and constitutively expressing PSA mutants

To distinguish whether the PSA promoter ‘OFF’ state that resulted from exposure to bacteriophages was due to induction or selection, we studied the interaction of Barc2635 with two mutants of *B. fragilis*: Δ*psa*^44^ and Δ*mpiM44*^43^ The former mutant lacks the PSA biosynthesis locus, and the latter has the PSA promoter locked in the ‘ON’ state, and therefore, constitutively synthesizes PSA. *In vitro* phage infection assays revealed comparable infection efficacy **[FIGURE S4C]** and no competitive advantage of either of the mutants during exposure to the bacteriophage **[FIGURE S4D-F]**. In agreement to the *in vitro* results, there was no selectivity of the bacteriophage towards either of the mutants in mice that were initially colonized with equal ratios of the two mutants **[FIGURE 4F, G]**. To note, the Δ*psa* mutant exhibited higher fitness in colonizing GF mice, however, this advantage remained unchanged regardless of exposure to the Barc2635 **[FIGURE S4G-H]**. This was further confirmed by comparing the ratio of Δ*psa/*Δ*mpiM44* on Day 10 to the ratio on Day 0 between the treated group and the control group **[FIGURE 4G]**. These data show that mutants lacking PSA do not have a fitness advantage during exposure to Barc2635.

### Bacteriophage-driven phase variation in *B. fragilis* results in reduction of Treg cells

It is well-established that the PSA of *B. fragilis* NCTC 9343 induces regulatory T cells (Tregs, CD4^+^Foxp3^+^RORgt^+^) in the colonic lamina propria of mice^13,45–47^. To monitor the effect of Barc2635 during *B. fragilis* colonization to the CD4^+^Foxp3^+^RORgt^+^ Tregs population, we extracted cells from the colonic lamina propria and immunophenotyped them by flow cytometry **[Figure 5A]**. We found that in the presence of Barc2635, the CD4^+^Foxp3^+^RORgt^+^ Tregs population is decreased concomitant with the decrease in *upaY* transcription, coinciding with less surface production of the immunomodulatory PSA **[Figure 5B**, **Figure 5C]**.

**Figure 5:**
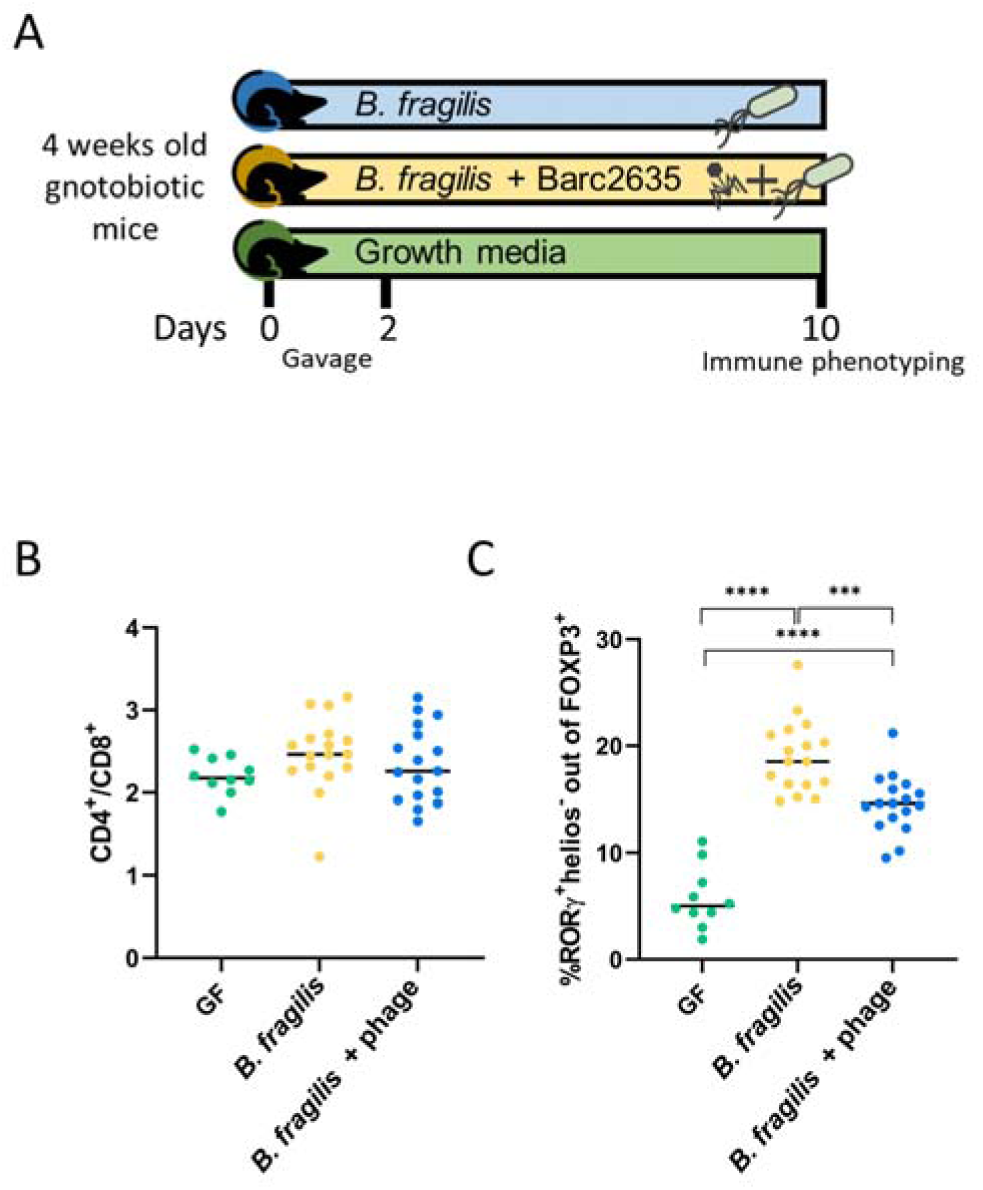
Bacteriophage-driven phase variation in B. fragilis results in reduction of Treg cells. **A**. Experimental design of *in vivo* experiments. **B.** CD4^+^ to CD8^+^ ratio out of CD45^+^TCR ^+^ live cells in colon. Single cells were isolated from colon lamina propria. Immune cells were analyzed by flow cytometry. Each dot represents a mouse. **C.** ROR t^+^ helios^-^ percentages out of FOXP3^+^ cells in colon. Single cells were isolated from colon lamina propria. Immune cells were analyzed by flow cytometry. Each dot represents a mouse. \*\*\**P*<0.001, \*\*\*\**P*<0.0001, one-way analysis of variance (ANOVA).

**Figure S5:**
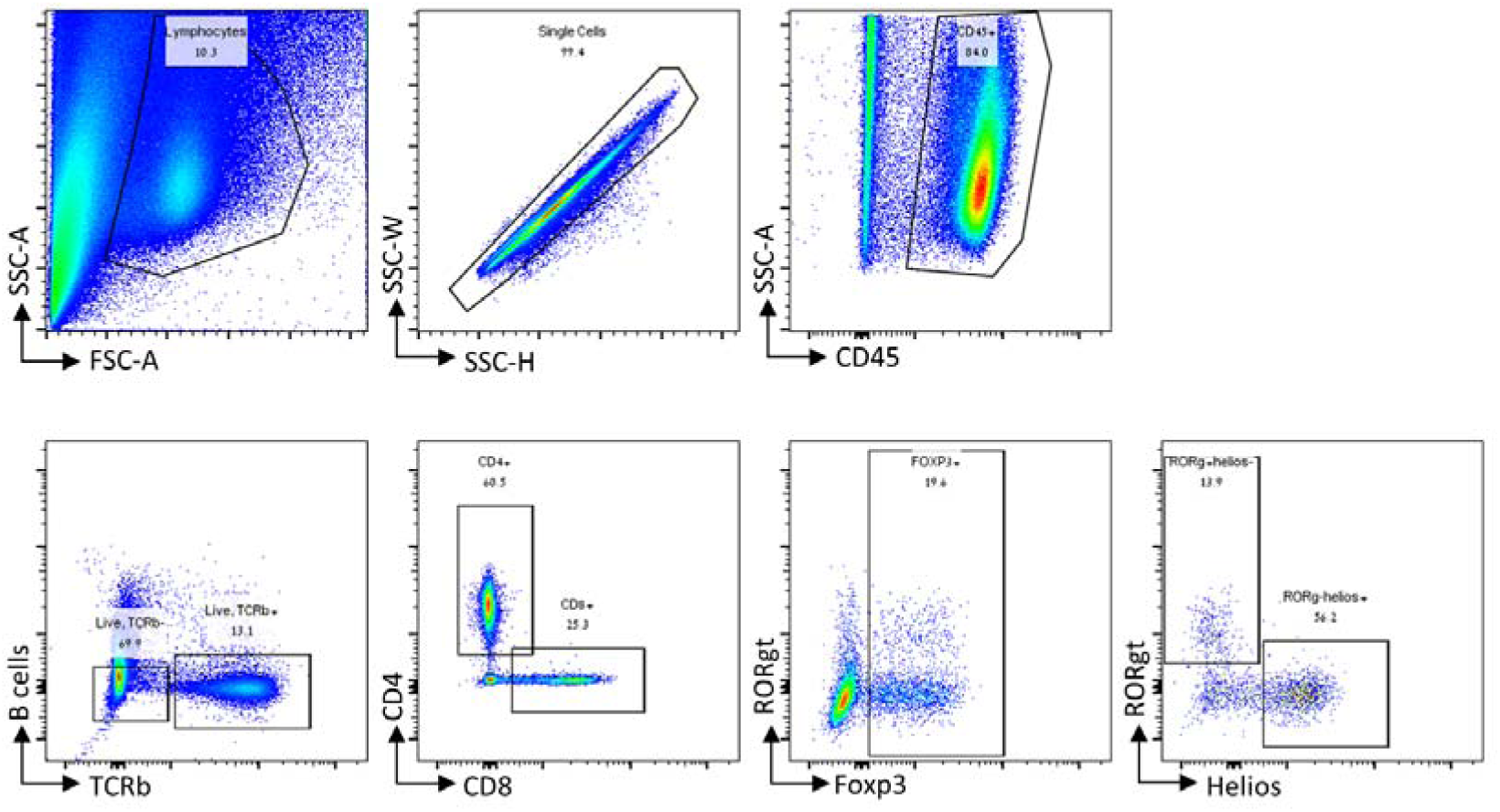
Gating strategy. Representative flow cytometry plots demonstrating the gating strategy for the staining panel.

## Discussion

Phase variations, prevalent in host-associated species, especially the gut Bacteroidales^9^, contribute to bacterial fitness in changing ecosystems. Reversible DNA inversions lead to phase variable synthesis of numerous molecules (e.g. surface, regulatory, and other molecules), and as such, confer functional plasticity. Our study reveals that the phase variable states of certain molecules correlate with gut inflammation, with potential implications on host physiology. By analyzing six different human gut metagenomic datasets with samples from IBD patients and healthy individuals,and using controlled mouse experiments, we identified multiple invertible genomic regions that map to 25 different Bacteroidales strains that are differentially oriented in disease and health. Notably, we find that both intergenic regions (e.g. promoters) and intragenic regions (e.g. genomic “shufflons”) exhibit altered orientations in gut inflammatory conditions – affecting gene expression.

The most prevalent genomic regions with differential orientations during inflammation and health were in polysaccharide (PS) promoters and near *susC/susD* homologs – both with immunomodulatory potential. SusC-like proteins are abundant β-barrel outer membrane proteins involved in nutrient acquisition in Bacteroidetes. Recently, an epitope of SusC proteins was shown to elicit T cell responses in IBD patients and healthy controls^48^, suggesting that operons that include SusC homologs might confer immunomodulatory properties to the bacteria. Among the phase-variable PSs, the anti-inflammatory PSA promoter of *B. fragilis* showed a higher percentage of reverse oriented reads in IBD patients compared to healthy controls, indicating that the ‘OFF’ orientation was more prevalent in IBD patients, potentially limiting the protective, anti-inflammatory, effects of PSA. These results align with a previous study^49^ that focused on the PSA promoter of *B. fragilis* in IBD patients using PCR digestion of biopsy samples. We further find that some DNA inversion states, like the PSA promoter of *B. fragilis*, are dynamic and can respond to changes in their local environment, specifically during inflammation.

Intriguingly, the preferential ‘OFF’ orientation of the PSA promoter of *B. fragilis* did not occur to the same extent in inflamed monocolonized mice as it did in inflamed mice colonized with a human microbiota and when bacteria were exposed to patients’ fecal filtrates. These results suggest that environmental factors in inflamed humanized mice and IBD patients are necessary for this effect. Our analysis of phage OTUs in the IBDMDB cohorts revealed enrichment of *B. fragilis* associated phage OTUs in samples where the PSA promoter was present in the ‘OFF’ orientation. In addition, exposure of *B. fragilis* to an isolated bacteriophage, Barc2635, resulted in higher frequency of the PSA promoter ‘OFF’ orientation, suggesting a role for gut bacteriophages in bacterial phase variation.

Our analyses were not able to distinguish the cause of the change in DNA orientation during inflammation and phage exposure. It is possible that these conditions induce inversions to the ‘OFF’ orientation but may also result from selection of bacteria that are not expressing PSA. Comparison of mutants that are unable or that constitutively synthesize PSA revealed similar relative PSA promoter orientations after Barc2635 exposure, suggesting that the observed decrease in PSA expression may be the result of induction of inversion rather than selection pressures. If so, this area is ripe for future mechanistic understanding.

The anti-inflammatory effects of PSA are mediated by upregulation of induced Tregs (Foxp3+RORgt+) and subsequent induction of IL-10 secretion^12,13^. We found that the Treg cells populations decreased in *B. fragilis* monocolonized mice infected with Barc2635, suggesting that the increase in bacteria with the PSA promoter in its’ ‘OFF’ orientation has implications on the host immune system, linking bacteriophage exposure to alterations in bacterial functionality with concomitant effects on host physiology. Applying the same analysis to *B. thetaiotaomicron*’s CPS3, we observed bacteriophages that are more abundant in samples with a higher ratio of CPS3 ‘ON’ oriented promoters, however, the bacteriophage to host abundances ratios per sample were too low to conclude that *B. thetaiotaomicron* bacteriophages lead to populations with altered CPS3 synthesis.

Altogether, we demonstrate differential orientations of Bacteroidales invertible regions in human IBD and identify bacteriophages as a potential trigger. These phase variations result in bacterial functional plasticity affecting the host immune system, correlating with inflammation. Bacteriophages are one factor that alter bacterial phase variable states. However, within the inflamed gut, there are numerous factors that may potentially alter the invertible states of Bacteroidales populations. These include the microbiome^35,39,50–52^ (e.g. neighboring microbes), the gut metabolome^50^, an altered immune system^53^ and physical alterations in the intestine such as abnormal pH concentrations^54^, osmotic^55,56^ and oxidative stress^57,58^. For example, Tropini et al.^59^ showed that PEG induced osmotic perturbation resulted in an increased ratio of CPS4 to CPS5 in *B. thetaiotaomicron*.

This study demonstrates that alterations in invertible genomic regions, and consequently molecular phase variations in gut bacteria during IBD, influence the host immune system and inflammatory state. These phase variations can occur in response to changes in gut environmental factors during inflammation, including, but not limited to bacteriophages. Future studies integrating bacterial DNA inversions and phase variation analyses may illuminate the role of bacterial functional plasticity in additional physiological states, possibly resulting from different environmental factors, which may drive alterations in microbe-host interactions.

### Limitations of the Study

The six datasets analyzed in this study included adults and children, healthy controls and inflamed patients (before and after treatment) and longitudinal samples. The impact of each of these factors, and the impact of diet and lifestyle, merit further research. The PhaseFinder^9^ algorithm that we applied identifies DNA inversions in short reads. As such, our analysis does not include other phase variation mechanisms like duplications or insertions. Also, analysis of short reads of metagenomics sequencing data may overlook inversions resulting from long structural genomic alterations^3,7,60^. Furthermore, metagenomic sequencing data averages the whole bacterial population of samples, implying a dilution factor for low abundant bacteria. Moreover, genomic inversion sites might interfere with genome sequence assemblies, leading to incomplete genomes, and thus could be overlooked when applying tools that search inversions in reference genomes. Also, our bacteriophage computational analysis, which is based on Nishiyama *et al.*^39^, might have overlooked some of the bacteriophages in the patients, since it relies on prior analysis of MAGs that were predicted as temperate phages. This study used an identified and isolated lytic bacteriophage for *B. fragilis*, further studies are required to elucidate the functional effects of IBD associated lytic and temperate bacteriophages. With these limitations notwithstanding, our study highlights the importance of considering bacterial phase variation in the context of IBD and its potential impact on inflammation and on altered microbe-host interactions.

## Methods

### Bacterial strains and growth conditions

*B. fragilis* NCTC 9343 was grown in Brain heart infusion (BHIS) supplemented with 5 mg/L hemin (Alfa Aesar) in 1 N NaOH, and 2.5 μg/L vitamin K under strictly anaerobic conditions (80% N2, 10% H2, 10% CO2) at 37 °C in an anaerobic chamber.

### Mice

All mouse work was in accordance with protocols approved by the local IACUC committee under approval numbers: IL-151-10-21 and IL-105-06-21.

4-5 weeks-old Germ-Free (GF) C57BL/6 mice (males and females) from the Technion colony were used. Mice were housed and maintained in a GF care facility and were provided with food and water *ad libitum*; they were exposed to a 12:12 h light-dark cycle at room temperature.

Humanized mice were generated by oral gavage of GF C57BL/6 mice with 200 µl mix of human fecal microbiota and cultured *B. fragilis* NCTC 9343. To prepare this mix, human feces from a single healthy human donor were suspended in sterile PBS 1:10 w/v (1 gram feces was suspended in 10 ml of sterile PBS). Next, 10^8^ CFUs of *B. fragilis* NCTC 9343 were resuspended in 1 ml of sterile PBS and added to 1 ml of fecal supernatant. *B. fragilis* NCTC 9343 were grown on Brain Heart Infusion agar plates (BHI, BD BBLTM) supplemented with 5 µg/ml hemin (Alfa Aesar) in 1 N NaOH and 2.5 µg/ml vitamin K (Thermo Fisher Scientific) in 100% EtOH, at 37 °C in an anaerobic chamber, 85% N2, 10% CO2, 5% H2 (COY). The progeny of these mice were co-housed, randomized, and grouped before treatment.

Gnotobiotic mice monocolonized with *B. fragilis* were created by oral gavage of GF C57BL/6 mice with 200 μl of 5×10^8^ CFU\ml of *B. fragilis* NCTC 9343, grown in the same conditions as above. The mice were co-housed, randomized, and grouped before treatment. Both gavages, in humanized and monocolonized mice, were performed once.

For phage immunomodulation studies, GF mice were gavaged twice (on day 0 and on day 2) with *B. fragilis* NCTC 9343 (10^8^ CFU in 200 µl Bacteroides phage recovery medium (BPRM) media) and bacteriophage Barc2635 (10^9^ PFU in 200 µl 0.22-µm-filtered BPRM), *B. fragilis* only (10^8^ CFU in 200 µl BPRM media), or growth media as control. On day 10 mice were sacrificed for bacteria and immune phenotyping.

### DSS model

For DSS experiments, male and female mice were co-housed separately. Mice were randomized and grouped (water control and DSS separately) before treatment. Throughout the experiments, close attention was paid to monitor any distress responses exhibited by the mice: mice with a 20% decrease in body weight, immobility, or an extreme reaction to touch, were excluded from the experiments.

For acute DSS-induced colitis in humanized mice, progeny of humanized mice were treated with 3% DSS in their drinking water for 9 days followed by 5 days recovery before sacrificing for the experiment. The control animals were administered distilled water.

Fecal samples were obtained on days 0, 6, 9 and 14 (before DSS treatment, during and after discontinuing DSS treatments), for further analyses.

For acute DSS-induced colitis in gnotobiotic mice colonized with *B. fragilis*, monocolonized mice were treated with 3% DSS in their drinking water for 7 days. The control animals were administered distilled water.

Fecal samples were obtained on days 0 and 7 (before and after DSS treatment), for further analyses.

### Mice Inflammation measurement

Fecal supernatants were prepared after suspension in 1:10 sterile PBS and centrifugation at 4,500×g for 15 minutes. Enzyme-linked immunosorbent assay (ELISA) to measure calprotectin concentrations was performed using Mouse S100A8/S100A9 Heterodimer kit according to the manufacturer protocol [R&D systems].

Mice weights were assessed using an electronic scale at the same day-time.

### qPCR and primers

DNA was extracted from fecal samples (30-50 mg) using ZymoBIOMICS DNA Miniprep Kit [Zymo research]. The ‘ON’/’OFF’ orientation of the PSA promoter in the extracted DNA samples was determined by quantitative polymerase chain reaction (qPCR) using SYBR® Green mix [Thermo Fisher Scientific]. Two sets of primers were designed to target the PSA locus **[Figure S1A-C]**. One set targeted the reference gene *upaY*, the first gene immediately downstream to the promoter region, and used as a proxy to the number of bacteria in the samples. The second set of primers targeted the promoter region and would only produce a product when the orientation is ‘ON’. The ratio of ‘ON’/’OFF’ PSA orientation in the samples was calculated using the 2^−ΔΔCT^ method^61^ calculated against the PSA locked ‘ON’ results (100% ‘ON’ orientation).

*upaY* gene expression was determined by RT-qPCR with the *upaY* primers. RNA was extracted from fecal samples using zymoBIOMICS RNA miniprep kit [Zymo research]. Following DNAse treatment (NEB), reverse transcription of RNA to cDNA was performed using the qScript cDNA Synthesis Kit [Quantabio]. *upaY* mRNA levels were determined by qPCR using SYBR® Green mix [Thermo Fisher Scientific] with primers against *rpsL* as a reference gene. Controls for non-template and non-reverse transcription were included. Primer efficiency was determined for each set of primers^62^. The 2−ΔΔCT method was employed for the specificity fold change tests.

Primers:

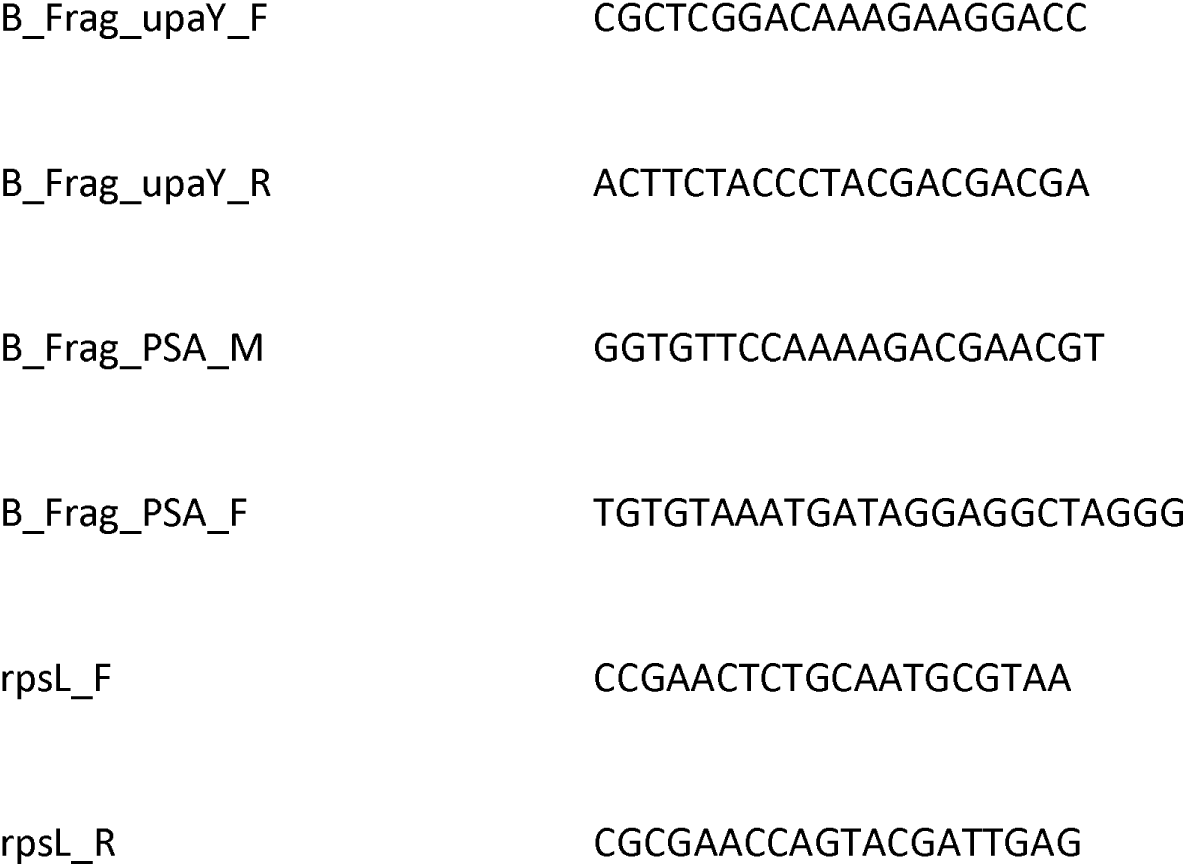

### Mice feces sequencing

DNA was extracted from fecal samples (30-50 mg) using ZymoBIOMICS DNA Miniprep Kit [Zymo research]. Samples underwent quality control by Qubit fluorescence analysis to determine the concentration of DNA for downstream analysis (ThermoFisher, Cat. Q32850). Libraries were prepared using the Illumina Tagmentation DNA prep streamlined library preparation protocol according to the manufacturer’s instructions with a minimum of 50 ng of DNA starting mass and 8 cycles of PCR enrichment, ending with a fragment size of 550 bp. IDT for Illumina DNA/RNA UD indexes and Nextera DNA CD indexes were used (Illumina IDT, Cat. 20027213; Illumina Nextera, Cat. 20018708).

All libraries were diluted to 15 pM in 96-plex pools and validated on 100-cycle paired-ends read Miseq V2 runs (Illumina, Cat. MS-102-2002), before shipping to the US at 4 nM for sequencing on the Novaseq 6000 in S4 mode at 96-plex in a 300-cycle paired-end reads run, with an estimated read depth of 30 Gbp per sample (Illumina, Cat. 20028312). Final loading concentration of 600 pM. All sequencing runs were performed with a spike-in of 1% PhiX control library V3 (Illumina, Cat. FC-110-3001). The sequencing mean library size was 134,629,540.5 reads [range: 10,107,679 – 396,239,822].

### Taxonomic profiling

Community profiling was performed using metaphlan4^63^ v4.0.059 with mpa database vJan21. For each sample, the forward reads were first aligned against the mpa database using bowtie2 v2.3.5.160 (flags **`--sam-no-hd –-sam-no-sq –-no-unal –-very-sensitive`**). Next, the resulting sam file was analyzed by metaphlan4 with default parameters, resulting in a merged relative abundances table.

### Microbiome analysis

Initial visual exploration of sequenced data was conducted using the MicrobiomeAnalyst^64^ web platform and followed by a comprehensive statistical analysis of using the Phyloseq^65^ version 1.44.0 and vegan^66^ version 2.6-4 packages in R v4.3.1. Read counts were transformed into relative abundances by normalization to the total number of reads per sample. Low-abundance filters were applied to discard species whose relative abundance did not reach 0.1% and did not appear in at least 20% of the samples. Alpha diversity (observed species and Shannon diversity) was calculated at the species level using the ‘plot_richness’ function and compared with the Kruskal–Wallis rank sum test. Beta diversity distance matrices (Aitchison distance) were ordinated using the ‘ordinate’ function, compared using the vegan^66^ package’s ‘adonis’ function (Permutational Multivariate Analysis of Variance), and visualized using PCoA. Beta dispersion (Aitchison distance) was calculated using the vegan^66^ package’s ‘betadisper’ function. Differential abundance of species between mice groups and timepoints were computed with R package Maaslin 2^67^, version 1.14.1 with CLR transformation of the data. Plotting was performed using R packages tidyr version 1.3.0, reshape2 version 1.4.4, ggpubr version 0.6.0, ggplot2^68^ version 3.4.3, and EnhancedVolcano version 1.18.0.

### Analysis of differential abundances of phages from Nishiyama *et al.*^39^

Count tables, metadata, and the sequences of MAGs and viral regions from Nishiyama *et al.*^39^ analysis of the IBDMDB dataest were obtained from the following site: ftp://ftp.genome.jp/pub/db/community/ibd-phage/.

Statistical analysis was done using the Phyloseq^65^ version 1.44.0 and DESeq2^69^ version 1.40.2 version 2.6-4 packages in R v4.3.1. The data was filtered to include only viral OTUs, and Low-abundance filters were applied to discard OTUs whose had less than 10 read counts and did not appear in at least 20% of the samples. Read counts were normalized to relative abundances. Samples were split to low (<40%) or high (>60%) ratios of the PSA promoter ‘ON’ orientation, according to the PhaseFinder results. Differential abundance of viral OTUs between the groups were computed with R package DESeq2^69^ with default settings. Plotting was performed using R packages tidyr version 1.3.0, reshape2 version 1.4.4, ggpubr version 0.6.0, ggplot2 version 3.4.3, and EnhancedVolcano version 1.18.0. Phage-to-host abundance ratio was calculated for each phage-*B. fragilis* and phage-*B. thetaiotaomicron* pairs according to the bacterial host predictions done by Nishiyama *et al.*^39^.

### Identification of DNA inversion regions

Representative Bacteroides species were selected from using the following approach: database of human-associated microbial species was constructed from 118K metagenomics assembled genomes (MAGs) recovered from human-associated metagenomics samples acquired from Pasolli *et al.*^70^, Almeida *et al.*^71^, and Forster *et al.*^72^. Taxonomy assignment was performed using gtdbtk v2.0.0 and GTDB release 207 with the classify_wf program and default parameters^73^. Overall, 34 *Bacteroides* species were identified of which 33 had species-level GTDB taxonomy assignments. For each of these species, the representative genome from GTDB was used for the PhaseFinder analysis. We have also included the genomes of three additional *B. fragilis* strains, *Phocaeicola dorei*, and *Parabacteroides distasonis*. Overall, 39 genomes were included. Information about the genomes is provided in Table S1.

PhaseFinder^9^ (v1.0) was used to identify invertible regions in metagenomics samples. The default parameters of PhaseFinder were used. Putative inversion regions were detected by identifying inverted repeats in bacteroidales genomes using the ‘locate’ function. A database containing the inversion regions forward and reverse orientations was created using the tool’s ‘create’ function. Using the tool’s ‘ratio’ function, metagenomic samples from the publicly available cohorts were then aligned to the database, resulting in the ratio of reverse oriented reads out of all reads assigned to each region. We filtered the results by removing identified regions with <20 reads supporting either the forward or reverse orientations combined from the paired-end method, mean Pe_ratio <1% across all samples, and sites within coding regions of rRNA products. To compare DNA orientation patterns between CD and UC patients with those of healthy controls, we focused on sites that displayed a difference of over 10% between at least one of the groups. The Wilcoxon rank sum test was employed to conduct these comparisons, and the Benjamini-Hochberg method was utilized to correct for multiple comparisons, with a false discovery rate (FDR) set at less than 0.1. Each invertible region was manually curated to assess its coding regions, gene annotations, and their putative functions. Briefly, genomic regions were visualized online using the NCBI Graphical Sequence Viewer (version 3.47.0). Invertible regions with coding sequences (CDS) within them, were annotated according to the CDS name(s). Invertible regions lacking CDS within them were searched for CDSs that start in proximity to the invertible DNA sites (<200∼bp). CDSs in the region (four upstream and four downstream) were used to assess the functionality of the region. Regions containing or in proximity to rRNA and tRNA genes were filtered from the comparisons as well as invertible regions with no CDSs start in proximity to the inverted repeats.

### Bacteriophage isolation

Bacteriophage BARC2635 was isolated from raw sewage. Briefly, inflowing raw sewage for a waste-water treatment plant (WWTP) from Barcelona (Spain) was filtered through low protein binding 0.22 µm pore size polyethersulfone (PES) membrane filters (Millex-GP, Millipore, Bedford, Massachusetts) to remove bacteria. Isolated plaques were obtained by the double-agar layer technique^74^. Briefly, tubes containing 2.5 ml of soft BPRM–agar kept at 45°C were inoculated with 1 ml of an exponential growth phase culture (OD600=0.3, corresponding to ca 2×10^8^ CFU / ml) of the host bacteria grown in BPRM broth and 1 ml of the filtered sewage sample. After gently mixing, the contents of each tube were poured onto a plate of BRPM-agar and incubated inside GasPak (BBL) jars at 37°C. Plaques were clearly spotted after 18 h of incubation.

For phage isolation, discrete well-isolated plaques were stabbed with a sterile needle and inoculated in a tube containing 5 ml of BRPM broth. Then 1 ml of a culture of *B. fragilis* NCTC 9343 in exponential growth was inoculated into the tube, which was then incubated for 18h at 37 °C. After incubation, an aliquot of the culture was treated with chloroform (1:10 (v;v), vigorously mixed for 5 minutes and centrifuged at 16,000×g for 5 minutes^75^. The supernatant containing the phage suspensions were further filtered through low protein binding 0.22 µm pore size polyethersulfone (PES) membrane filters (Millex-GP, Millipore, Bedford, Massachusetts), diluted and plated as indicated in the previous paragraph to verify the uniformity of the plaques. Then, one well differentiated plaque was stabbed and the whole operation was repeated to obtain a high titer, over 1×10^9^ plaque forming units (PFU), phage suspensions.

### Phage genome sequencing

Phage particles were PEG-precipitated (6,000-12,000 MW, 8%), and isolated by a CsCl gradient; 33 g, 41 g, 55 g in 50 ml TM buffer (50mM Tris-Cl pH8.0, 10mM MgCl2), ultracentrifuged at average 152,000×g for 1.5hrs, and dialyzed overnight in 100mM Tris ph7.5, 1M NaCl, 1mM EDTA^58^. Genomic DNA was extracted using phenol-chloroform as described^76^.

Illumina sequencing of Bacteroides phage Barc2635 was performed at the Biopolymers Facility, Harvard Medical School, Department of Genetics, producing paired-end reads of 150 bp. Adapter sequence removal and quality trimming was performed using BBDuk, part of the BBTools (v 37.50) suite of programs. The reads were further screened against NCBI’s UniVec_Core database (build 10.0) and the *B. fragilis* NCTC 9343 genome sequence using blastn and reads that returned a significant hit to either were removed. The phage genome was assembled de novo using Velvet 1.2.10 under a k-value determined by Velvet Optimizer (v. 2.2.5). The genome was annotated using a customized version of Prokka^77^ v1.12, altered to additionally utilize the profile Hidden Markov Model (HMM) libraries of Pfam^78^ version 35, TIGRFAM^79^ version 15, and the Clusters of Orthologous Genes^80^ (COGs, 2020 update, and PRotein K(c)lusters (PRK) portions of NCBI’s Conserved Domain Database^81^ during annotation., submitted to NCBI, and assigned GenBank accession MN078104. Phage Genome map **[Figure S3C]** was visualized using the online tool Proksee (https://proksee.ca/, accessed on 21 January 2023).

### Barc2635 susceptibility and competition assays

Frozen stocks of *B. fragilis* NCTC 9343 Δ*mpiM44*^43^ (genetically engineered bacteria that constitutively express PSA) and *B. fragilis* NCTC 9343 Δ*PSA*^44^ (lacks PSA biosynthesis locus) maintained in 25% glycerol at −80 °C were thawed on Brain Heart Infusion agar plates (BHI, BD BBLTM) supplemented with 5 µg/ml hemin (Alfa Aesar) in 1 N NaOH and 2.5 µg/ml vitamin K (Thermo Fisher Scientific) in 100% EtOH, at 37 °C in an anaerobic chamber, 85% N2, 10% CO2, 5% H2 (COY). Then strains were grown anaerobically for up to 3 days, and a single colony was picked for each bacterial strain, inoculated into 5 ml BPRM and grown anaerobically overnight to provide the starting culture for experiments. A dilution of 1:10 was done the following day and the diluted culture was incubated at the same conditions until mid-logarithmic phase of growth.

At the beginning of the experiment, comparable CFU amounts of each strain were verified at OD 0.5 (Δ*mpiM44* 2×10^8^ and Δ*PSA* 2×10^8^ ).

For susceptibility assays: 300µl of each strain was infected with Barc2635 in 1:100 ratio, and then incubated for 5 minutes in 37 °C, followed by a centrifuge of 4,500g x 5 minutes and re-suspension with clean 300µl BPRM to get rid of free phages that did not infect the cells. The infected bacterial cells were added to 3 ml of the molten soft top agar and mixed well before being poured onto the bottom agar. The following day plaques were counted according to the formula:

PFU/ml = # of plaques / (dilution * infection volume in ml).

For competition assays *in vitro*: Both bacterial strains (*ΔmpiM44* and Δ*PSA)* were grown at the same anaerobic conditions until mid-logarithmic phase of growth. Equal CFU’s were verified. Next, bacterial cells were mixed in 1:1 ratio and then Barc2635 was added to the mix at 1:100 ratio. 200µl of Time point (Tp) 0 (starting point of the experiment) and Tp2 (2 hours after Barc2635 infection) were taken to qPCR analysis.

In vivo competition was done by gavaging C57BL/6 mice with equal ratios of Δ*mpiM44* and Δ*psa*,wait two days for the bacteria to settle in and then Barc2635 (10^9^ PFU in 200 µl 0.22-µm-filtered BPRM) was added on Tp0. 10 days after the inoculation mice fecal samples were collected for qPCR analysis.

Primers used:

For Δ*mpiM44*

wcf_F: GGC CTC CTT CAT CTC AGG TTT ATC C

wcf_R: GAT AAT CGC GGC ACC CTA TGG G

For Δ*PSA*

mpi_F: AAG AGG GCT ATG TGT TTC AGG ACG

mpi_R: CTG CGT GCG AGA GCT TCT TTG

### Viral OTUs multiple alignment

Alignments and phylogenetic tree of whole genomes of viral OTUs identified as bacteriophages against *B. fragilis*^39^ as well as bacteriophage Barc2635 were generated using MAFFT^82^ online service (version 7). Multiple sequence alignment was done using the default settings of the site. An average linkage UPGMA guide tree was constructed using average-linkage UPGMA with the online MAFFT^82^ service and was visualized with R package ggtree^83^ (version 3.4.4).

### Fecal filtrates of patients

Recruitment of IBD patients for this study was conducted at the Rambam Health Care Campus (RHCC). The study was approved by the local institutional review boards with study numbers 0052-17 and 0075-09 in which all patients consented to be included in it. Fecal samples were collected by the patients, prior to their clinic visit. The samples were then stored at –80°C until they were shipped to the laboratory for analysis.

Calprotectin levels were measured in each fecal sample using LIAISON Calprotectin (catalog No. 318960) according to the manufacturer’s instructions. The levels of calprotectin in the fecal samples were used as a measure of disease activity in IBD patients.

### *In vitro* assays with fecal filtrates from IBD patients

*B. fragilis* NCTC 9343 was grown in Brain heart infusion (BHIS) supplemented with 5 mg/L hemin (Alfa Aesar) in 1 N NaOH, and 2.5 μg/L vitamin K to OD_600_∼0.6 and centrifuged at 4,500×g for 5 minutes. Bacterial pellets were washed twice with sterile PBS to remove BHIS components and then suspended with 1 mL supplemented M9 minimal media.

Patients’ fecal samples were suspended in sterile PBS (1:5), centrifuged at 4,500×g for 15 minutes, and supernatants were collected and filtered using the Medical Millex-VV Syringe Filter Unit, 0.22 µm, PVDF membrane.

For the *in vitro* assay, *B. fragilis* was cultured in a mixture of M9 and patients’ fecal supernatants in a 1:25:25 ratio (bacteria suspended in M9: M9: fecal supernatants) and grown at 37°C in an anaerobic chamber, 85% N2, 10% CO2, 5% H2 (COY). As a control, *B. fragilis* was grown in M9 and mixed with the same ratios but with sterile PBS instead of fecal filtrates. Subsequently, 200μl of each culture was collected for DNA extraction at OD_600_∼0.6 using ZymoBIOMICS DNA Miniprep Kit [Zymo research]. qPCR analysis was done using the primers used in mice experiments, with 3 technical replicates from each individual patient:

B_Frag_upaY_F CGCTCGGACAAAGAAGGACC

B_Frag_upaY_R ACTTCTACCCTACGACGACGA

B_Frag_PSA_M GGTGTTCCAAAAGACGAACGT

B_Frag_PSA_F TGTGTAAATGATAGGAGGCTAGGG

Each sample was technically repeated 3 times.

M9 Medium was prepared as follows:

18.7 mM NH_4_Cl

42.2 mM Na_2_HPO_4_

22 mM KH_2_PO_4_

8.5 mM NaCl

Supplements:

0.1 mM CaCl_2_.2H_2_O

1 mM MgSO_4_.7H_2_O

0.5% glucose

0.05% L-cysteine

5 g/L Hemin

2.5 mg/ml VitK_1_

2 mg/ml FeSO_4_.7H_2_O

5 ng/ml VitB12

### Gut lamina propria preparation

For lamina propria immunophenotyping, mice colons were removed by cutting the colon from the cecum-colon junction to the anus. Fat tissue was carefully removed from colon tissue and further processed for single cell suspension preparation using lamina propria dissociation kit (Miltenyi), according to the manufacturer’s protocol.

### Flow cytometry

Cell preparations for flow cytometry analysis were performed in 5 ml tubes or U shape 96 wells plates. Single cells were washed with PBS and stained for live/dead staining using 1:1,000 in PBS, Zombie fixable viability dye (Biolegend) for 10 minutes, at room temperature, and washed once with FACS buffer, by centrifuge at 300×g for 5 minutes. For FcR blocking, cells were incubated with 0.5 µg CD16/CD32 antibody for 10 minutes on ice and proceeded to further staining without a washing step. Extracellular markers were stained with the relevant antibody panels for 30 minutes on ice and washed twice with FACS buffer, by centrifuge at 300×g for 5 minutes. After the last wash, cells were fixed with Foxp3 Fixation/Permeabilization working solution (Thermo) for 16 hours at 4 °C in the dark. For Intracellular staining, cells were permeabilized using 1X Foxp3 permeabilization buffer (Thermo) according to the manufacturer protocol. For intracellular blocking, 2 µl of 2% rat serum (Stemcell technologies) was added to each well for 15 minutes at room temperature and proceeded to further staining without a washing step.

To quantify the percentage of *B. fragilis* from monocolonized mice with PSA on their surface, feces from monocolonized mice with *B. fragilis* with or without phage were suspended 1:10 in ice cold PBS (mg/µl) and centrifuged at 300×g for 5 minutes, 4°C. Supernatants were separated from pellets and further centrifuged at 4,500×g for 5 minutes, 4°C. Bacterial pellets were resuspended in an ice cold FACS buffer, 1:10 from initial PBS suspension. 100 µl of resuspended bacteria were incubated with 1:1,000 rabbit antibodies (Antibodies preparation previously described^1^) against *B. fragilis* PSA for 30 minutes at 4°C. Bacteria were washed twice using an ice cold FACS buffer by centrifuge at 4,500×g for 5 minutes and then incubated with a donkey anti rabbit fluorophore conjugated secondary antibody. After staining steps, the bacteria were washed twice with an ice cold FACS buffer and finally resuspended in 500 µl ice cold PBS plus 1:1,000 Hoechst dye and analyzed by flow cytometry using FSC and SSC thresholds of 1,000, and logarithmic scale. Gating strategy is detailed in **Figure S5**. Antibody specificity for PSA was verified using PSA mutant *B. fragilis.* Non-specific binding controls were included.

## Data availability

Sequencing data has been deposited in the NCBI sequence read archive (SRA) under the BioProject accession number: PRJNA916364.

## Acknowledgements

We would like to thank the Geva-Zatorsky lab members for fruitful discussions and contributions. Thanking Zachary Merenstein and Neekoo Farahmandpour for helping in establishing the qPCR method; Dr. Svetlana Friedman, Ran Tahan and Prof. Debbie Lindell for advising us on the bacteriophage characterization and Dr. Carolina Tropini for insightful discussions. We thank Drs. Amir Grau, Ofer Shenker, and the biomedical core facility at the Technion Rappaport faculty of Medicine for their help with flow cytometry; BiotaX labs for metagenomics sequencing, and Prof. Ogata and Dr. Nishiyama from the Institute for Chemical Research, Kyoto University, for helping in accessing their data. Figure 3A was created with BioRender.com.

## Author contributions

SC HH RZ DKK TG IS and NGZ conceived and designed the project. SC HH RZ DKK and TG performed and analyzed the experiments. SC, IS and NGZ planned the computational and analytical aspects; SC and IS performed the computational analyses. MJC and LC contributed to the computational analysis, phage sequencing and analysis, and interpretation of the results. SP and YC designed, led, and executed the clinical study. KJ contributed to the phage-bacteria-host study and to the bacteriophage morphological characterization. JJ isolated the bacteriophage. NM RN, and NBA helped with the experiments. SC HH RZ TG IS LC and NGZ wrote the manuscript. SC RZ HH and DKK contributed equally to the study. All authors read and approved the manuscript. NGZ conceived and planned the study, supervised it, interpreted the experiments, and wrote the manuscript.

