## Supplementary Figures S1-S5 for "Inflammation and bacteriophages affect DNA inversion states and functionality of the gut microbiota"

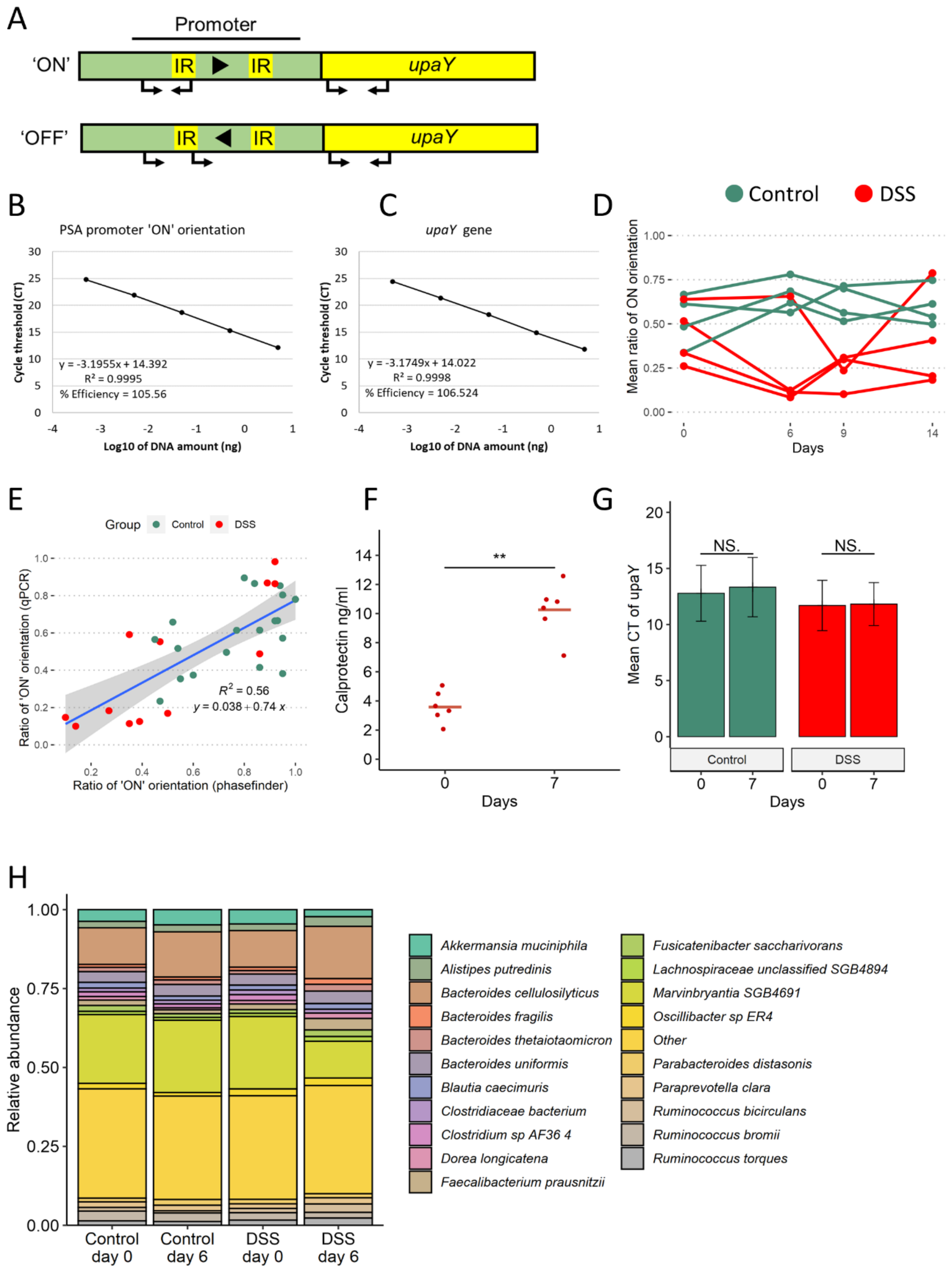


**FigureS1: Relative orientation of the PSA promoter of *B. fragilis* is affected by inflammation.**

**A.** Schematics of the qPCR assay used to assess the ratio of PSA 'ON' promoter orientation. Primers (arrows) were directed upstream and within the PSA promoter invertible region (IR: inverted repeats) targeting only the 'ON' oriented promoters. Another set of primers was directed at the *upaY* and used to normalize the results to the number of genomes in a sample. **B. & C.** Primer efficiencies for the two sets of primers. Standard curves were calculated for each primer set from the Cycle thresholds (CTs) and the log of the initial DNA concentrations. Each sample represents the mean CT of 3 technical repeats. **D.** Mean ratio of *B. fragilis* PSA's promoter 'ON' orientation measured by qPCR for different experiments on different days (n=8-12 in each time point). Green: Control group, Red: DSS treated mice. **E.** Scatter plot of the ratio of *B. fragilis* PSA's promoter 'ON' orientation measured by qPCR and by the PhaseFinder tool. Each point represents a single sample, colored according to group and timepoints: green: Control group, red: DSS treated mice. Blue line represents linear regression (gray area represents 95% confidence intervals). **F.** Calprotectin levels (ng/ml) measured in gnotobiotic mice colonized with *B. fragilis* and treated with DSS on different days of the experiment. Line represents the mean. (Wilcoxon rank sum test, $^{**}p < 0.01$) **G.** Mean qPCR cycle threshold (CTs) of *upaY* gene DNA, representing the abundance of *B. fragilis* in gnotobiotic mice monocolonized with *B. fragilis* of different groups and timepoints. Bars represent the means of the CTs in each group and timepoint. Error bars represent the standard error. Green: Control group, Red: DSS treated mice. (Wilcoxon rank sum test, $p > 0.05$). **H.** Stacked bar plots representing the relative abundance of the 20 most abundant bacterial species in the humanized mice in each experimental group in days 0 and 6.

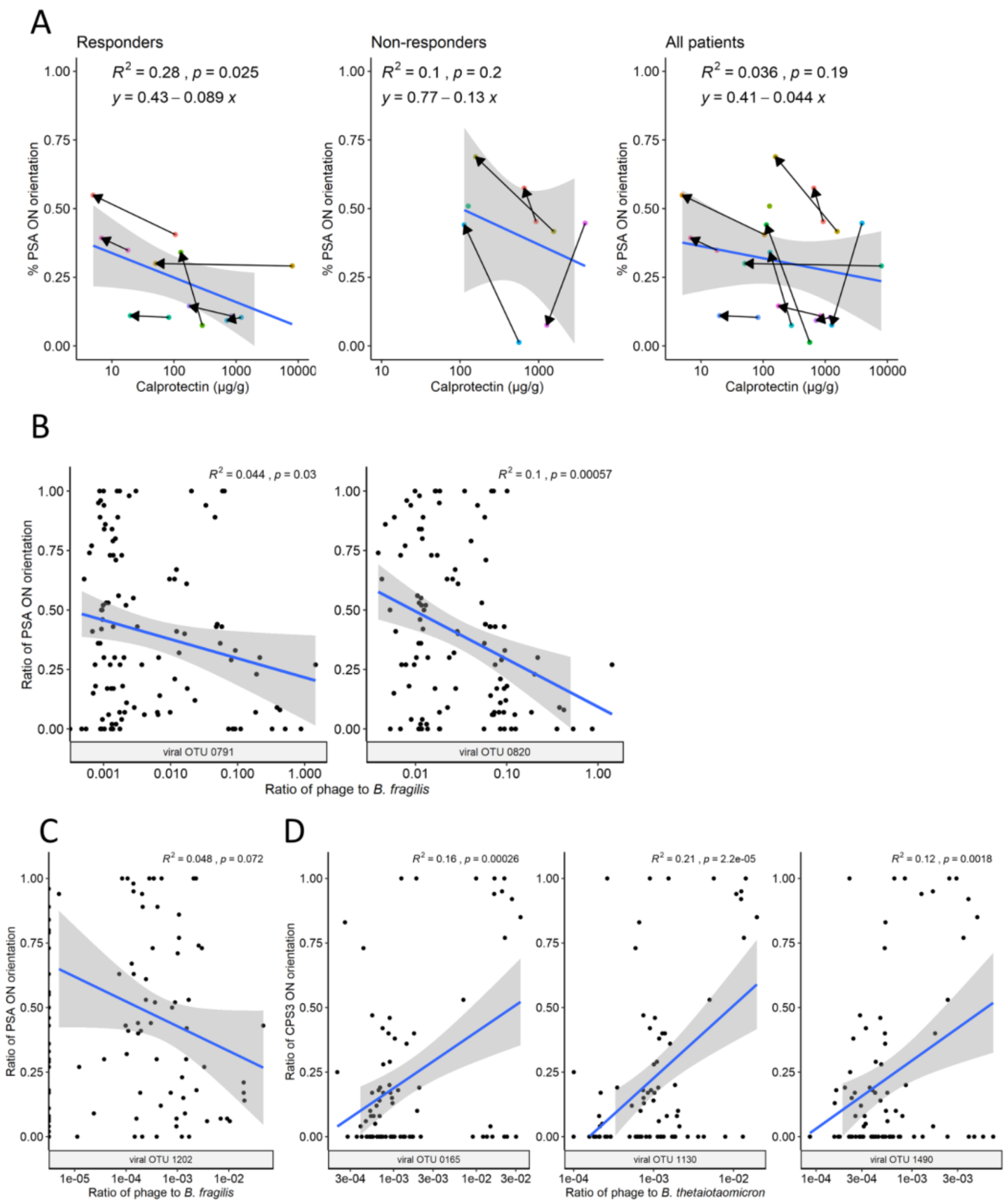


**Figure S2: Phage correlation with *B. fragilis* polysaccharide A promoter orientation.**

**A.** Scatter plots of the ratio of *B. fragilis* PSA's promoter 'ON' orientation measured by qPCR and fecal calprotectin levels (µg/g). Each point represents a single sample, colors indicate the same patient, arrows denote the change from samples before anti-TNF treatment to after. Blue line represents linear regression (gray area represents 95% confidence intervals). **B. & C.** Scatter plots of the ratio of *B. fragilis* PSA's promoter 'ON' orientation measured by qPCR and phage to host ratios of viral OTUs predicted to infect *B. fragilis* in Nishiyama *et al.* (2020). Blue line represents linear regression (gray area represents 95% confidence intervals). **D.** Scatter plots of the ratio of *B. fragilis* PSA's promoter 'ON' orientation measured by qPCR and phage to host ratios of viral OTUs predicted to infect *B. thetaiotaomicron* in Nishiyama *et al.* (2020). Blue line represents linear regression (gray area represents 95% confidence intervals).

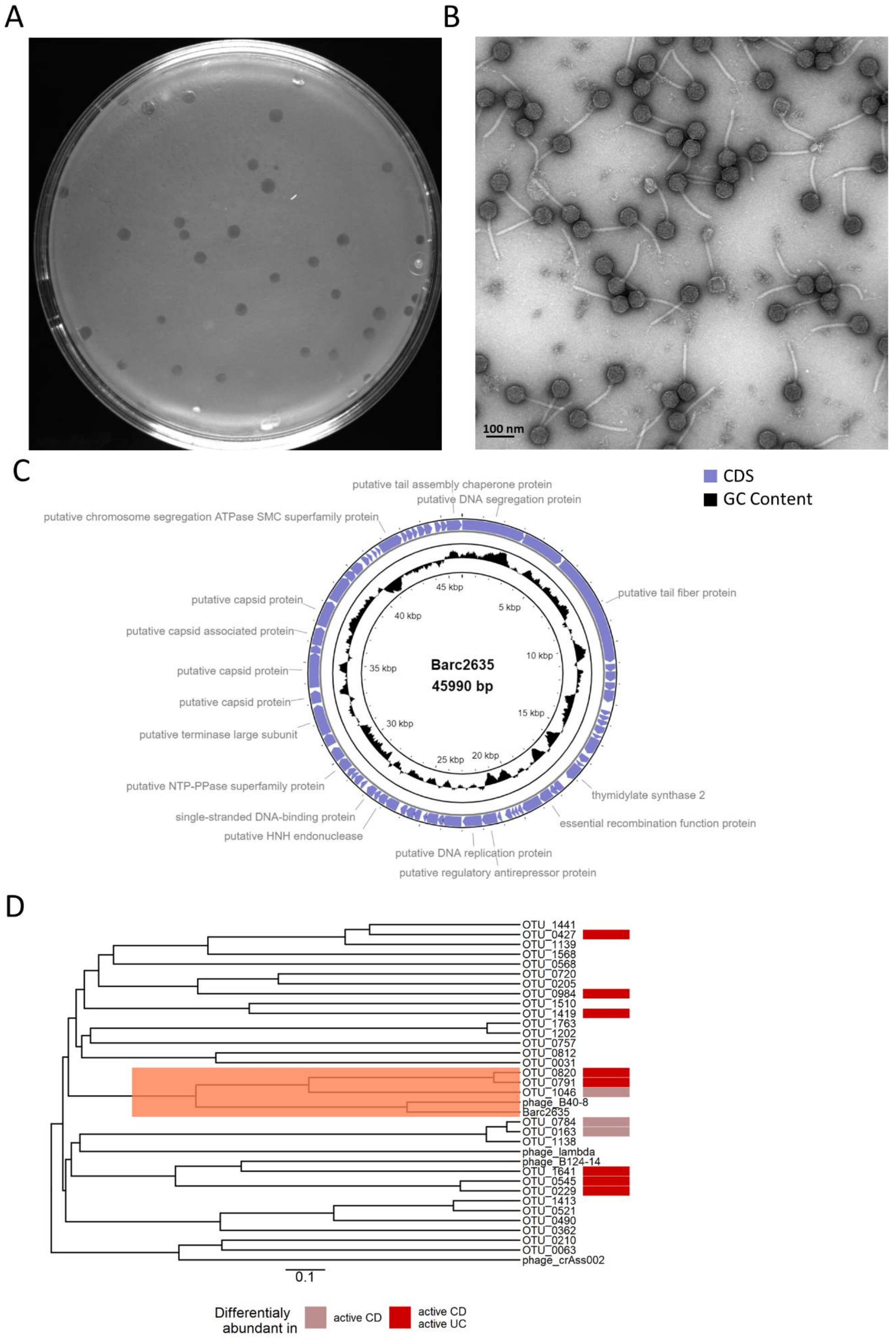


**Figure S3: Characterization of Bacteriophage Barc2635**

**A.** Barc2635 plaques on a lawn *B. fragilis* NCTC 9343. **B.** A representative transmission electron microscopy of Barc2635. **C.** Genome structure, GC content, and putative annotations of Barc2635. CDS: coding sequence. **D.** Phylogenetic tree based on the whole genome of viral OTUs identified as bacteriophages against *B. fragilis* as well as *Bacteroides* bacteriophages Barc2635, B40-8, B124-14, crAss002, and *Enterobacteria* phage lambda. MAFFT was used to perform multiple sequence alignment and the average-linkage method was used to construct the phylogenetic tree. Colors denote association between the viral OTUs abundances in Nishiyama *et al.* (2020), IBDMDB cohort, with active Crohn's disease (light red) or with both active Ulcerative colitis and active Crohn's disease (red).

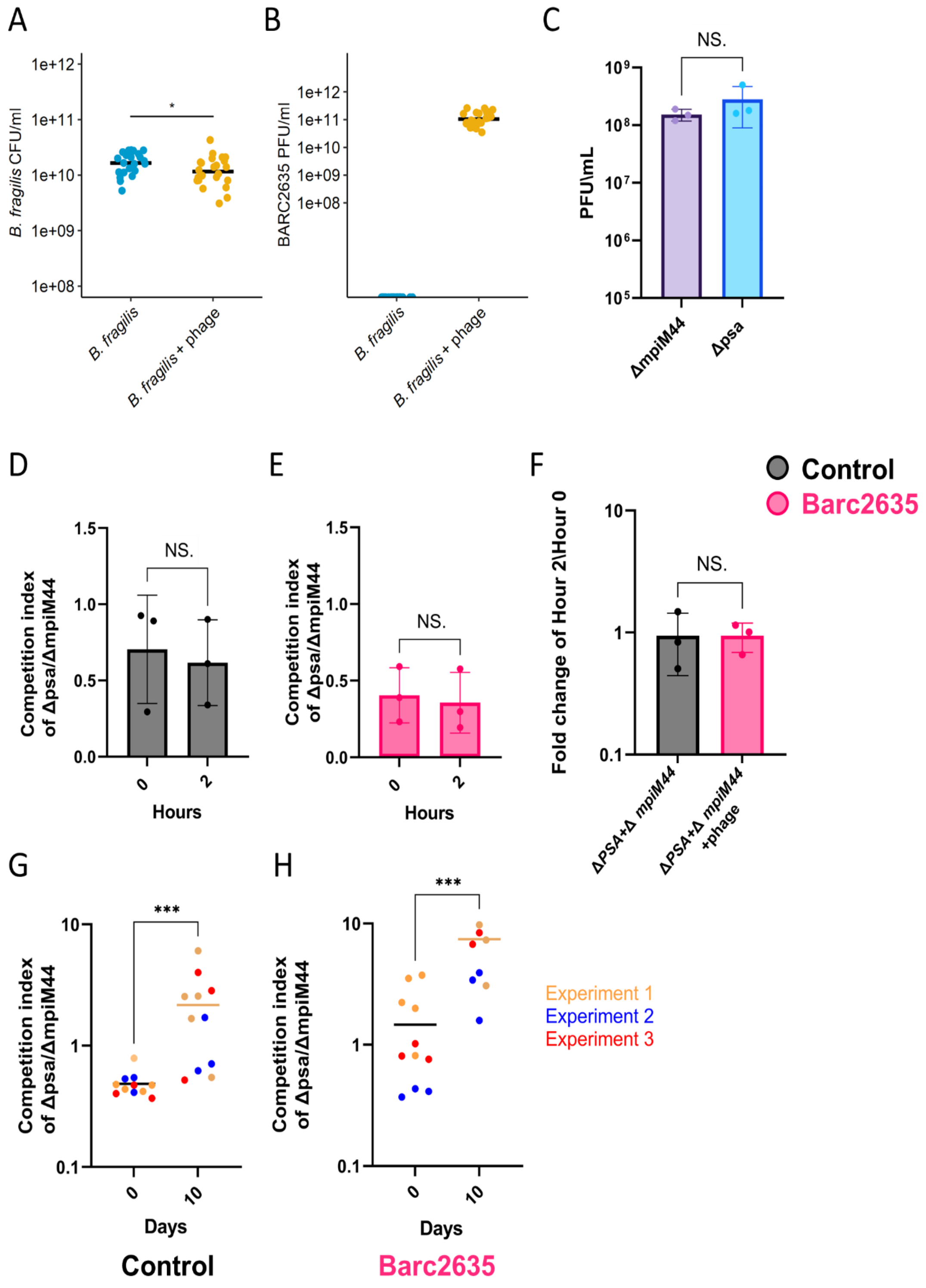


**FigureS4: Barc2635 infects both Δmpi and Δpsa in the same manner in vivo.**

**A.** CFUs of *B. fragilis* in fecal samples of mice monocolonized with *B. fragilis* (blue) or colonized with *B. fragilis* + phage (yellow) measured on day 10 (Wilcoxon rank sum test, *p < 0.05). **B**. PFUs of Barc2635 in fecal samples of mice monocolonized with *B. fragilis* (blue) or colonized with *B. fragilis* + phage (yellow) measured on day 10. **C**. Barc2635 plaque forming units of ΔmpiM44 (purple) compared to Δpsa (blue). Each dot represents a different experiment. (Mann-Whitney test, p>0.05). **D**. Competition index of Δpsa\ΔmpiM44 in the control group at the beginning of the experiment (0 horse) compared to 2 hours, measured by qPCR. Each dot represents a different experiment (Mann-Whitney test, p>0.05). E. Competition index of Δpsa\ΔmpiM44 in Barc2635 treated group at the beginning of the experiment (0 horse) compared to

2 hours, measured by qPCR. (Mann-Whitney test, $p>0.05$) **F.** Fold change of Δpsa\ΔmpiM44 at the beginning of the experiment (0 horse) compared to 2 hours in the Barc2635 bacteriophage treated group (pink) compared to Δpsa\ΔmpiM44 at Tp2 and Tp0 in the control group (black). Each dot represents a different experiment. (Mann-Whitney test, $p>0.05$). **G.** Competition index of Δpsa\ΔmpiM44 in the control group on day 0 compared to day 10, measured by qPCR. Each dot represents a mouse (Mann-Whitney test, ****p<0.001*). **H.** Competition index of Δpsa\ΔmpiM44 in the treated group on day 0 compared to day 10, measured by qPCR. Each dot represents a mouse (Mann-Whitney test, ****p<0.001*)

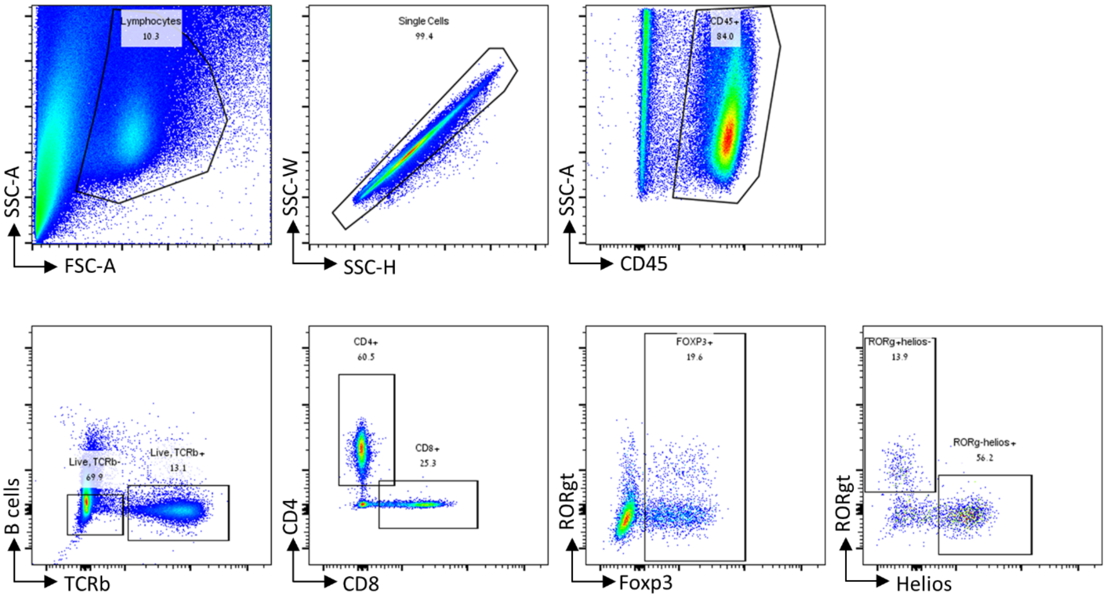


**Figure S5: Gating strategy.**

Representative flow cytometry plots demonstrating the gating strategy for the staining panel.
