## Supplementary figures and images for "Inflammation and bacteriophages affect DNA inversion states and functionality of the gut microbiota"

### Figure S1

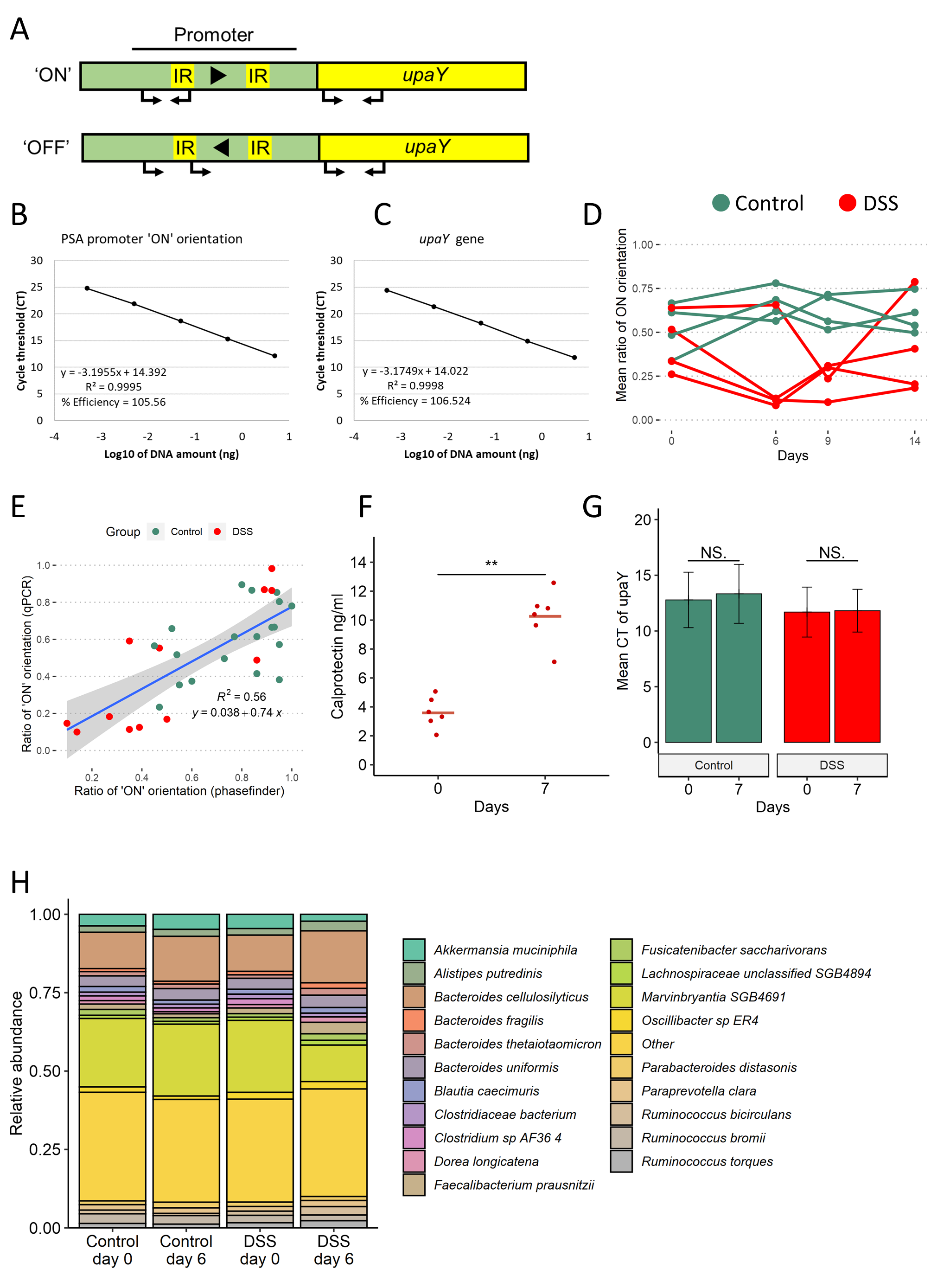

### Figure S2

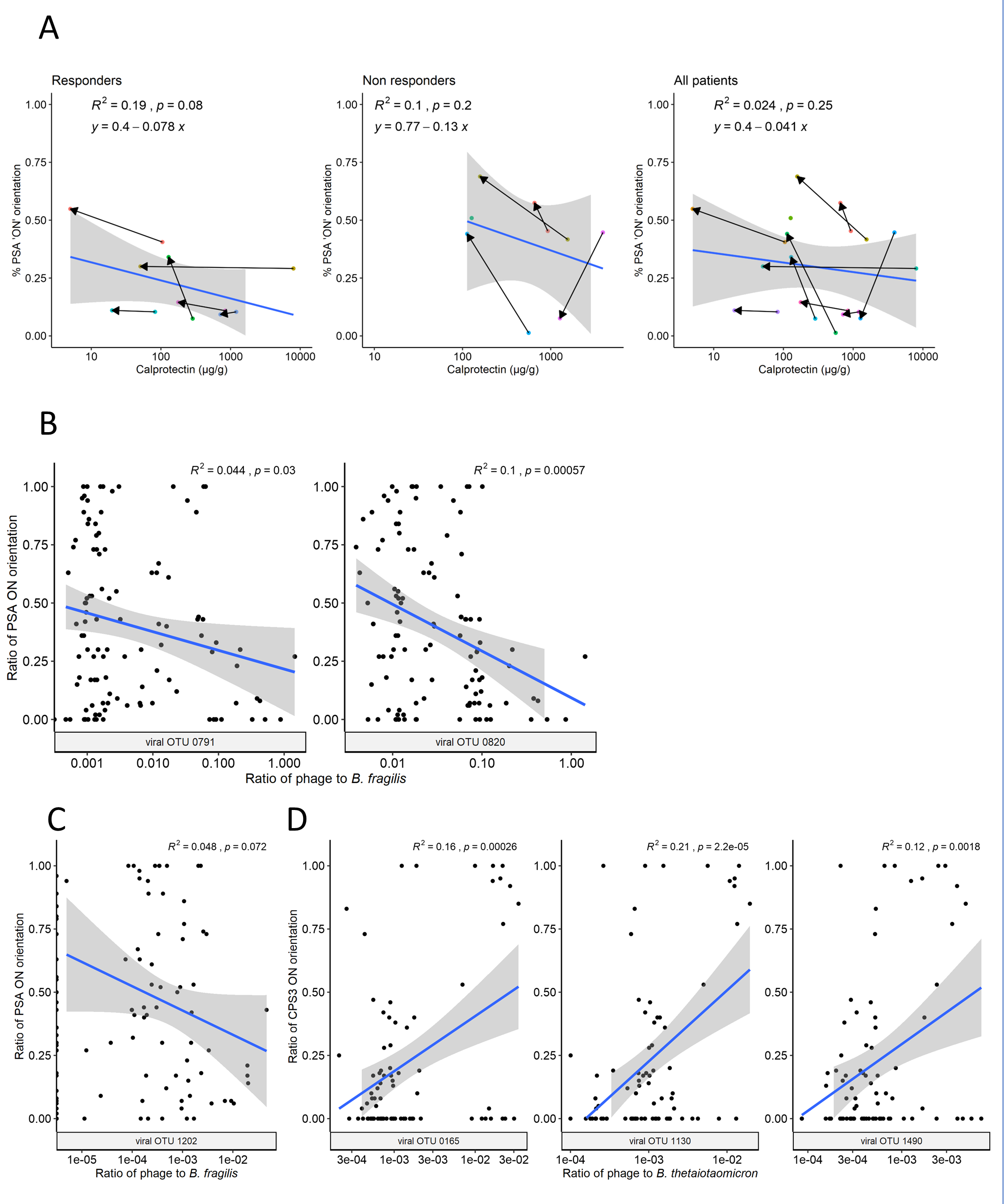

### Figure S3

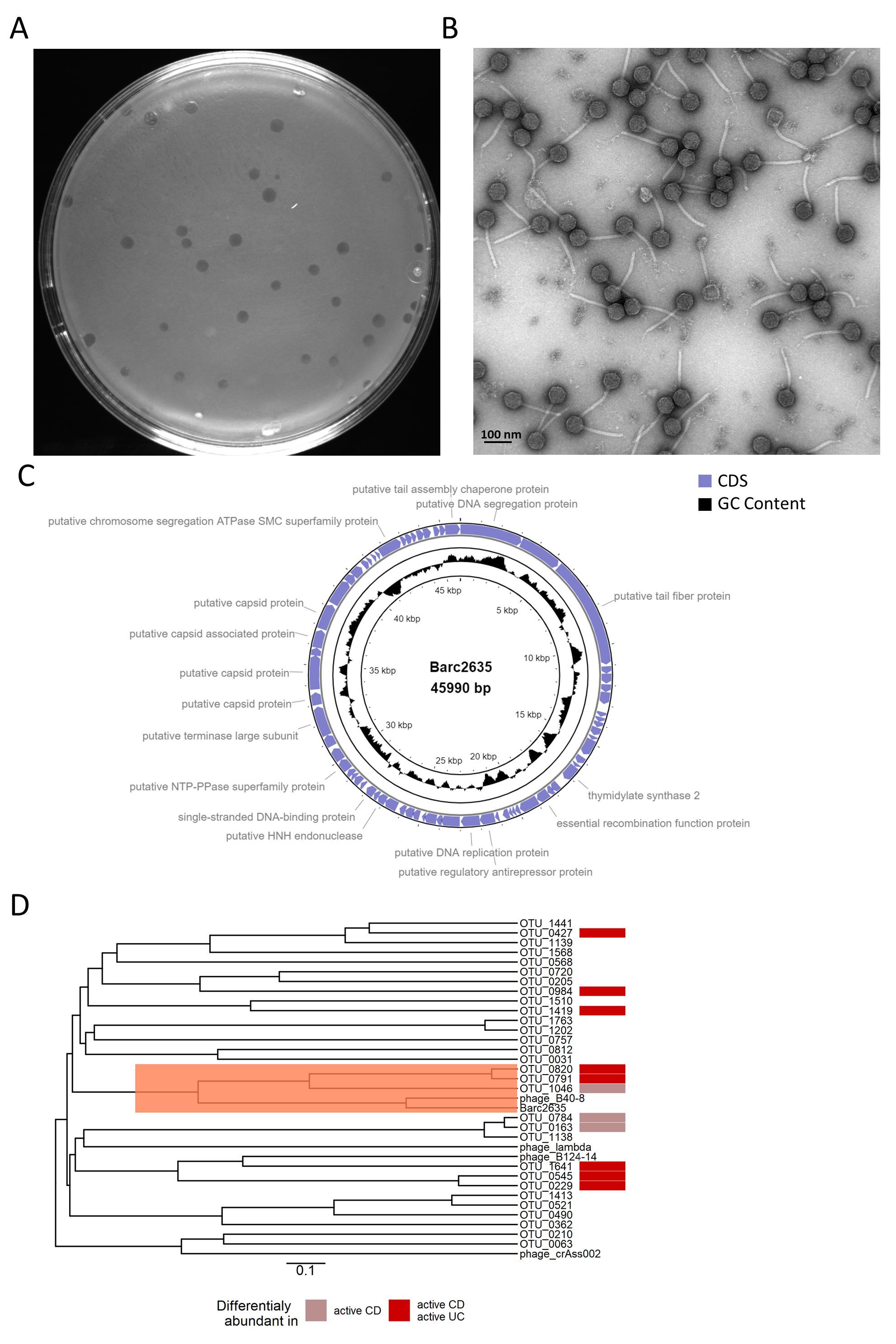

### Figure S4

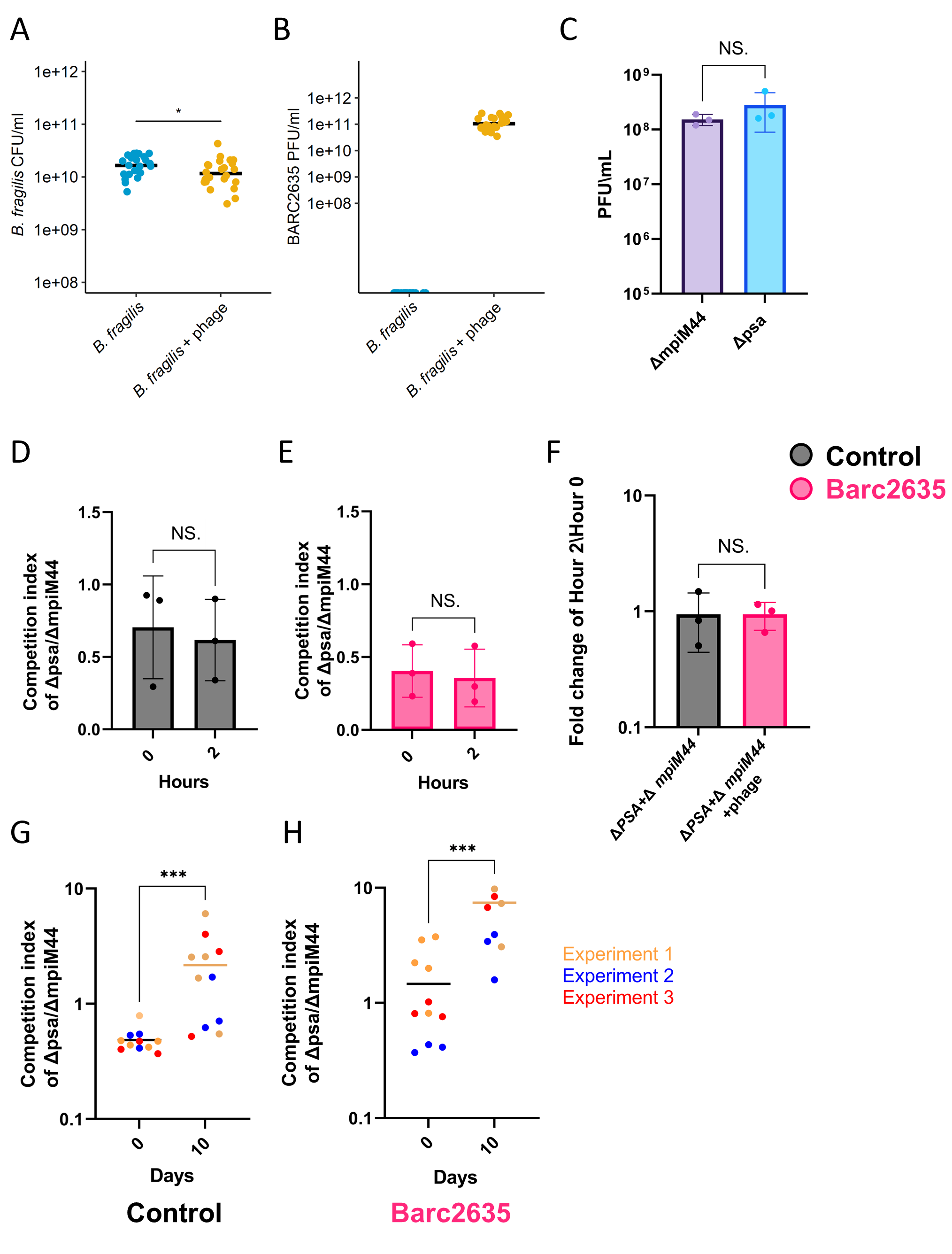

### Figure S5

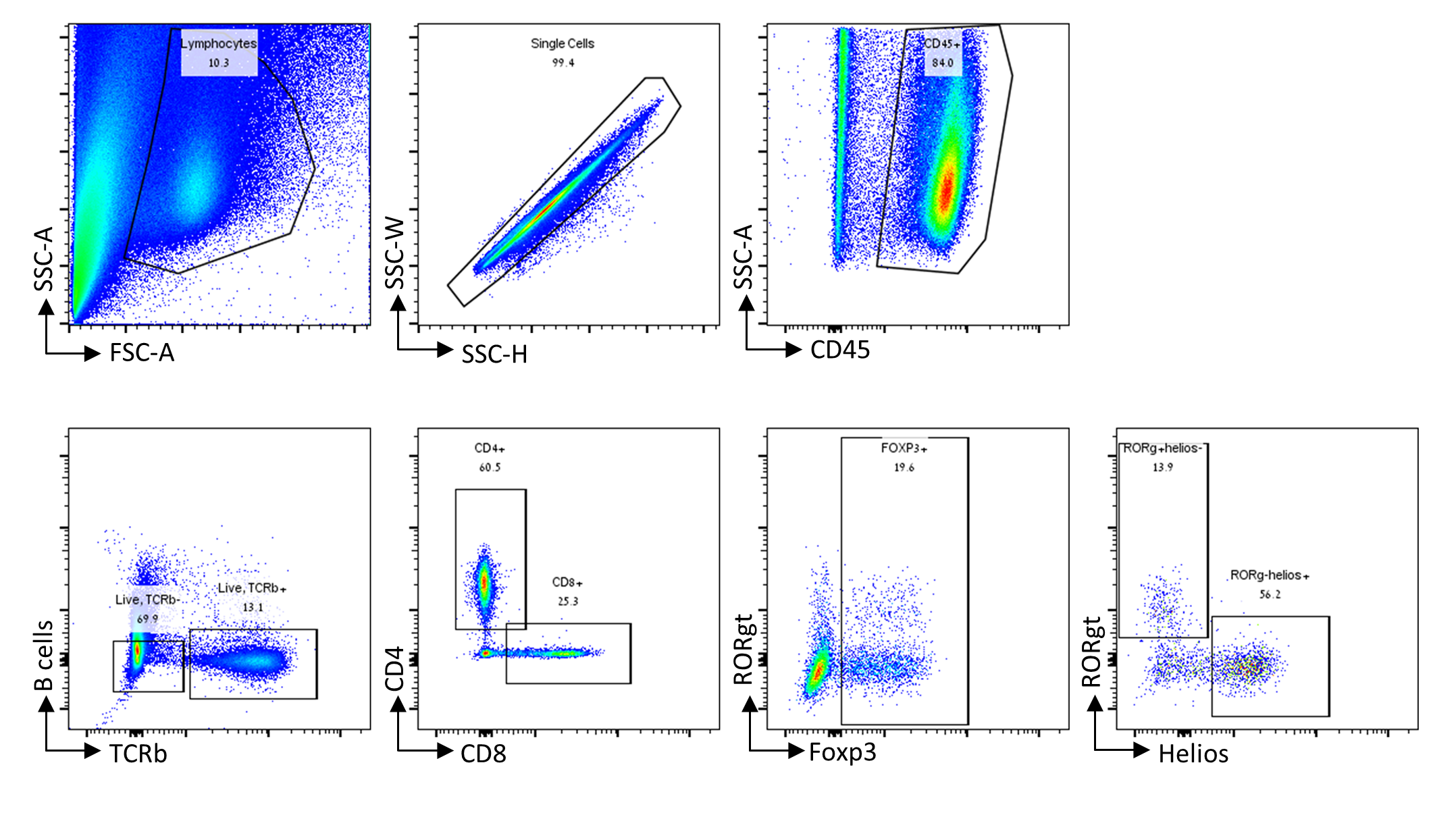
